# Introgression and Parental Conflict Shape Repeated Occurrences of Postzygotic Isolation

**DOI:** 10.1101/2025.08.12.669975

**Authors:** Megan E. Frayer, Hagar K. Soliman, Pia F. Schwarz, Jenn M. Coughlan

**Author notes:** These authors contributed equally to this manuscript (Megan E. Frayer, Hagar K. Soliman).

## Abstract

Postzygotic reproductive isolation is often thought to accumulate as a byproduct of neutral divergence. Yet, it frequently evolves rapidly, in line with non-neutral evolution. A major driver of intrinsic postzygotic reproductive barriers are intragenomic conflicts, such as conflict between maternal and paternal interests in resource allocation to offspring (i.e. parental conflict). Parental conflict may underlie hybrid seed inviability, a common and rapidly evolving reproductive barrier in angiosperms. Yet, in closely related, hybridizing species, it remains unclear how intragenomic conflicts and introgression interact to determine the fate of incompatibility alleles in nature. Here, we explore repeated incidences of hybrid seed inviability in a rising model: the *Mimulus guttatus* species complex. Using an extensive, range-wide crossing survey, we discover patterns of hybrid seed inviability within the widespread *M. guttatus* that are better described by geography than phylogeny. These patterns of reproductive isolation transgress species boundaries, as geographically-proximate but phylogenetically-distant species also exhibit similar patterns of hybrid seed inviability with allopatric populations of *M. guttatus*. We find strong support that patterns of reproductive isolation are consistent with parental conflict. Lastly, we provide evidence that introgression may underlie shared patterns of hybrid seed inviability between two species within this complex. Such introgression could have led to cascading reproductive isolation with other closely related species, creating a complex landscape of incompatibility. Overall, this work suggests that parental conflict and introgression can interact to shape the rapid and repeated evolution of strong reproductive isolation in the wild.

## Introduction

Understanding the origin and maintenance of species is a fundamental question in biology. Central to both of these processes is the emergence of reproductive isolation—the mechanisms that reduce interspecific fertilization and/or hybrid fitness. The evolution of intrinsic postzygotic isolation is often thought of as a slow accumulation of neutral or nearly neutral differences between two lineages (Coyne and Orr 1989, 1997, 2004). However, intrinsic postzygotic incompatibilities can also evolve rapidly and dynamically at various stages of divergence, suggesting a role of non-neutral processes in their evolution (Barbash et al. 2003; Johnson 2010; Corbett-Detig et al. 2013; Crespi and Nosil 2013; Coughlan and Matute 2020). Such rapid evolution of hybrid inviability and sterility has been observed in diverse species (Barbash et al. 2003; Charistianson et al. 2005; Briscoe Runquist et al. 2014; Schumer et al. 2014; İltaş et al. 2021; Gustafsson et al. 2022; Fusca et al. 2025). Thus, although not every species will rapidly accumulate intrinsic postzygotic isolation, this phenomenon can be found universally, providing important insights into how closely related taxa diverge in fundamental developmental processes, such as gametogenesis or offspring development (Charistianson et al. 2005; Wiley et al. 2009; Bracewell et al. 2011; Moyle et al. 2012; Brekke and Good 2014; Briscoe Runquist et al. 2014; Barnard-Kubow et al. 2016; Gustafsson et al. 2022).

One source of rapid evolution of incompatibilities are intragenomic conflicts, such as parental conflict in angiosperms and placental mammals (Haig and Westoby 1989, 1991; Moore and Haig 1991; Zeh and Zeh 2000; Burt and Trivers 2009; Coughlan 2023b; Soliman and Coughlan 2024; Frayer et al. 2025). Parental conflict occurs when unequal relatedness among offspring in a brood leads to opposing parental optima for resource allocation (Trivers 1974; Haig and Westoby 1989, 1991; Haig 1997). Under this theory, selection should favor paternal alleles that draw more resources from the maternal parent to their offspring, while maternal alleles arise to combat that excess via resource repression (Trivers 1974; Haig 1997). This parental “tug of war” can lead to an evolutionary arms race wherein sets of maternal and paternal alleles become coadapted to each other within a given population or species (Brandvain and Haig 2018). Parental conflict is thought to manifest most significantly in taxa that have evolved specialized tissues that facilitate nutrient exchange from maternal parents to the offspring, namely the endosperm in angiosperms and the placenta in eutherians (Soliman and Coughlan 2024). Mechanistically, such tug-of-war battles over resources are proposed to occur through genomic imprinting, a specialized form of gene regulation that has independently evolved in angiosperms and eutherians, wherein genes exhibit parent-of-origin biased expression (Haig and Westoby 1991).

Parental conflict theory posits that in crosses within a lineage, maternally- and paternally-acting alleles should be evenly matched, resulting in typical offspring development. However, crosses among lineages with different histories of conflict can reveal selfish alleles with such parent-of-origin effects, leading to dysfunctional offspring development and potentially death (Zeh and Zeh 2000; Gutierrez-Marcos et al. 2003; Coughlan 2023b,c). Several testable predictions arise from parental conflict theory: First, incompatibility should involve tissues that facilitate nutrient exchange among maternal parents and offspring (Soliman and Coughlan 2024). Second, reciprocal F1 offspring should exhibit parent-of-origin specific growth abnormalities. Namely, F1s sired by a lineage that has experienced a longer or more intense history of conflict should exhibit an overgrowth phenotype known as “paternal excess”, while the reciprocal cross should exhibit an undergrowth phenotype (i.e. “maternal excess”; Haig and Westoby 1991; Brandvain and Haig 2018). We refer to these as high- and low-conflict lineages, and note that these definitions align with high- and low-endosperm balance number, which is a similar conceptual framework more commonly used in the crop breeding literature (Johnston et al. 1980; Johnston and Hanneman 1982). Third, parental conflict theory suggests that any cross that restores the balance of maternally- and paternally-acting alleles should be compatible. As such, crosses between two lineages with similar histories of conflict should be compatible, even if their histories of conflict are evolutionarily independent (Coughlan et al. 2020). This creates a final testable prediction that the outcome of crosses should be predictable based on inferred histories of conflict.

In angiosperms, overgrowth phenotypes often involve delayed cell division or cellularization in the endosperm, both of which tend to be lethal (Rebernig et al. 2015; Oneal et al. 2016; Lafon-Placette et al. 2017, 2018; Roth et al. 2018; Coughlan et al. 2020; Sandstedt and Sweigart 2022). Conversely, undergrowth phenotypes often involve smaller seeds caused by accelerated endosperm development (Rebernig et al. 2015; Oneal et al. 2016; Lafon-Placette et al. 2018; Coughlan et al. 2020; İltaş et al. 2021; Sandstedt and Sweigart 2022). Hybrid seed inviability can evolve rapidly (Coughlan 2023b) including in taxa as recently diverged as 10,000 years ago (İltaş et al. 2021). Importantly, hybrid seed inviability is common—it has been reported in diverse angiosperm species (Grant 1966; Briscoe Runquist et al. 2014; Rebernig et al. 2015; Wolff et al. 2015; Florez-Rueda et al. 2016; Oneal et al. 2016; Lafon-Placette et al. 2017, 2018; Roth et al. 2019; Coughlan et al. 2020; Sandstedt et al. 2020; Ostevik et al. 2021; Sandstedt and Sweigart 2022; Butel et al. 2024). The observation that hybrid seed inviability has evolved rapidly and independently across angiosperms, with consistently similar patterns of dysfunctional seed development, indicates that it is highly repeatable, at least at the phenotypic level. However, it remains unclear how repeatable this barrier is at a genetic level or, if incompatibility alleles are shared, what the source of this shared genetic variation is. These questions have fundamental implications for evolution, as they can reveal how much of the genome is susceptible to genetic conflict and how quickly these alleles can arise and spread.

Introgression is a common way for alleles to spread among populations (Taylor and Larson 2019; Dagilis et al. 2022). While typically thought to hinder speciation, introgression can also contribute to reproductive isolation by introducing novel ecologically-mediated reproductive barriers (Long and Rieseberg 2024; Rosser et al. 2024). Introgression from one species into another can also enable reproductive isolation with a third species (Moran et al. 2024; Aguillon et al. 2025). In the case of incompatibility alleles evolving under intragenomic conflicts, theory predicts that once a selfish allele invades a new population, it should spread (Meiklejohn et al. 2018; Wang et al. 2024). Evidence of introgressed drivers has been found in several species (Meiklejohn et al. 2018; Svedberg et al. 2021; Wang et al. 2024), including those involved in incompatibility (Widen et al. 2023). However, it is unclear under what conditions an intrinsic incompatibility that has evolved via intragenomic conflicts could persist as a reproductive barrier. Furthermore, if and how theory developed for elements such as meiotic drivers translates to an allele evolving via parental conflict remains unknown.

Here, we leverage a rising model system for ecology, evolution, and genetics—the *Mimulus guttatus* species complex—to understand the repeated occurrence of hybrid seed inviability. We identify several instances of hybrid seed inviability across geographically and/or phylogenetically distinct taxa. We test whether these patterns of crossability conform to the prediction of parental conflict. We then assess whether the genetic basis of hybrid seed inviability between two incidences is shared, and whether they have a history of introgression. Our results support the idea that introgression of selfish elements can lead to the repeated evolution of reproductive isolation between lineages.

## Results

### Repeated occurrence of hybrid seed inviability in the Mimulus guttatus species complex

The *Mimulus guttatus* species complex is a relatively young clade of monkeyflowers endemic to the western half of North America (Vickery 1974, 1978; Fig.1A,B; crown age estimate: 674kya, Sandstedt et al. 2020). Using a range-wide crossing survey of 23 populations of *M. guttatus* crossed in a nearly full diallel design (426/506 unique directions, 1,742 crosses), we identified strong and asymmetric patterns of hybrid seed inviability between geographically distinct populations of *M. guttatus* (Fig.1B,C; Fig.S1; Table S1). Crosses between populations from the Sierra mountains/foothills and coastal or more northern parts of the range were largely inviable, manifesting as flat seeds that did not germinate when the Sierran populations served as the sire. In contrast, the reciprocal crosses produced smaller, viable seeds with high germination rates (Fig.1C; Fig.S2; Sierran x Northern *M. guttatus*: χ^2^ = 1427, DF = 3, *p* < 0.0001; Sierran x Coastal *M. guttatus*: χ^2^ = 2724, DF = 3, *p* < 0.0001). There was little to no hybrid seed inviability observed in crosses between populations of *M. guttatus* from the north and coast (herein “Northern” and “Coastal”) or within the Sierran lines— these crosses largely formed plump, viable seeds with high germination rates (Fig.S2; Northern x Coastal *M. guttatus*: χ^2^ = 6.88, DF = 3, *p* = 0.07). Despite broad consistency, we found some population-specificity in the magnitude of hybrid seed inviability. The most variable patterns of hybrid seed inviability arose from crosses between Sierran *M. guttatus* and Northern *M. guttatus*, and were largely caused by two Northern *M. guttatus* accessions that exhibited more viable seeds with Sierran lines and fewer with other Northern lines (i.e. CRES and TOK; Fig.1C). Furthermore, the Sierran *M. guttatus* themselves do not form a monophyletic group (Fig.1A). This finding highlights a novel incidence of hybrid seed inviability segregating within geographically distinct populations of *M. guttatus*, albeit with some population-specificity in the magnitude of incompatibility.

**Figure 1.**
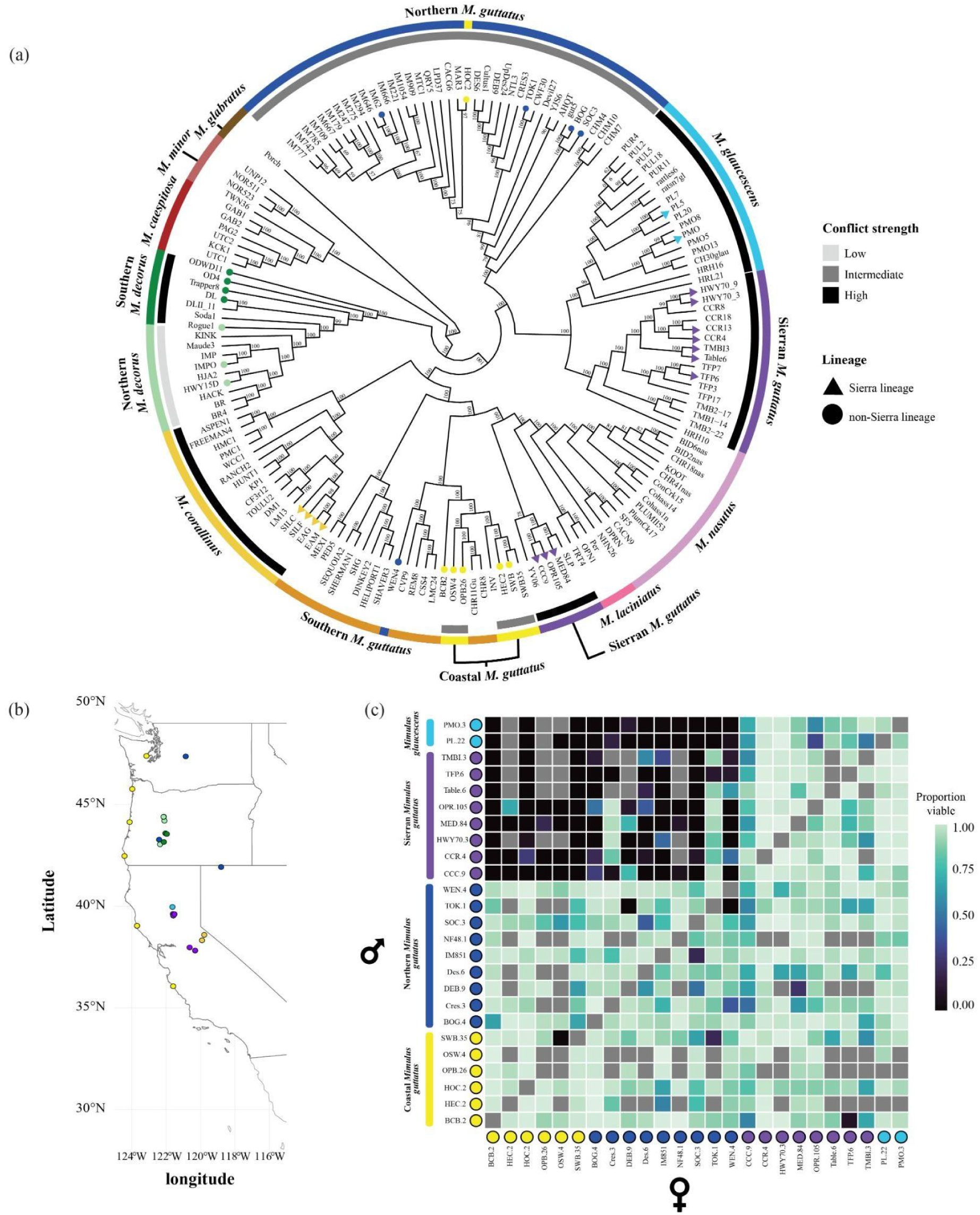
Hybrid seed inviability segregates within the widespread *M. guttatus* species complex. **A)** A maximum likelihood phylogeny of the *M. guttatus* species complex with lines used in the crossing survey denoted by colored shapes at the tips. Inferences of the history of conflict that a subset of these taxa have experienced are denoted by the inner greyscale ring. These inferences are based on the current work, as well as that of others (Coughlan et al. 2020). The outer colored ring denotes named species or lineages of *M. guttatus* and *M. decorus* that are inferred to have different histories of conflict. **B)** The geographic distribution of the lines used in the crossing survey. Samples are colored according to species/lineage. **C)** Proportion of viable seeds for pairwise crosses survey between each of the 25 populations. Grey boxes denote incomplete crosses.

Given the geographic pattern of incompatibility, we next tested whether two additional Sierran endemic species (*M. glaucescens* and *M. corallinus*) also exhibited this incompatibility. We crossed 2 populations of *M. glaucescens* to 21 populations of *M. guttatus* from our original crossing survey. Both Sierran *M. guttatus* and *M. glaucescens* behaved similarly in crosses with either Northern or Coastal *M. guttatus*, although seed inviability tended to be more severe in crosses sired by *M. glaucescens* (Fig.1). Although crosses between Sierran *M. guttatus* and *M. glaucescens* produced overall plump, viable seeds, we did observe a small but significant decline in viable seed production when *M. glaucescens* sired the cross (∼ 6-7%; χ^2^ = 36.4, DF = 3, *p* < 0.001; Fig.S2), consistent with previous work on sympatric populations of *M. glaucescens* and Sierran *M. guttatus* (Ivey et al. 2023). We then crossed another, more distantly related Sierran endemic, *M. corallinus* to a single population of Northern *M. guttatus.* We again found strong and asymmetric hybrid seed inviability, wherein seeds were inviable when *M. corallinus* sired these crosses but were viable when *M. corallinus* served as the dam (Fig.2B; χ ^2^ = 68, DF = 3, *p* < 0.0001).

**Figure 2.**
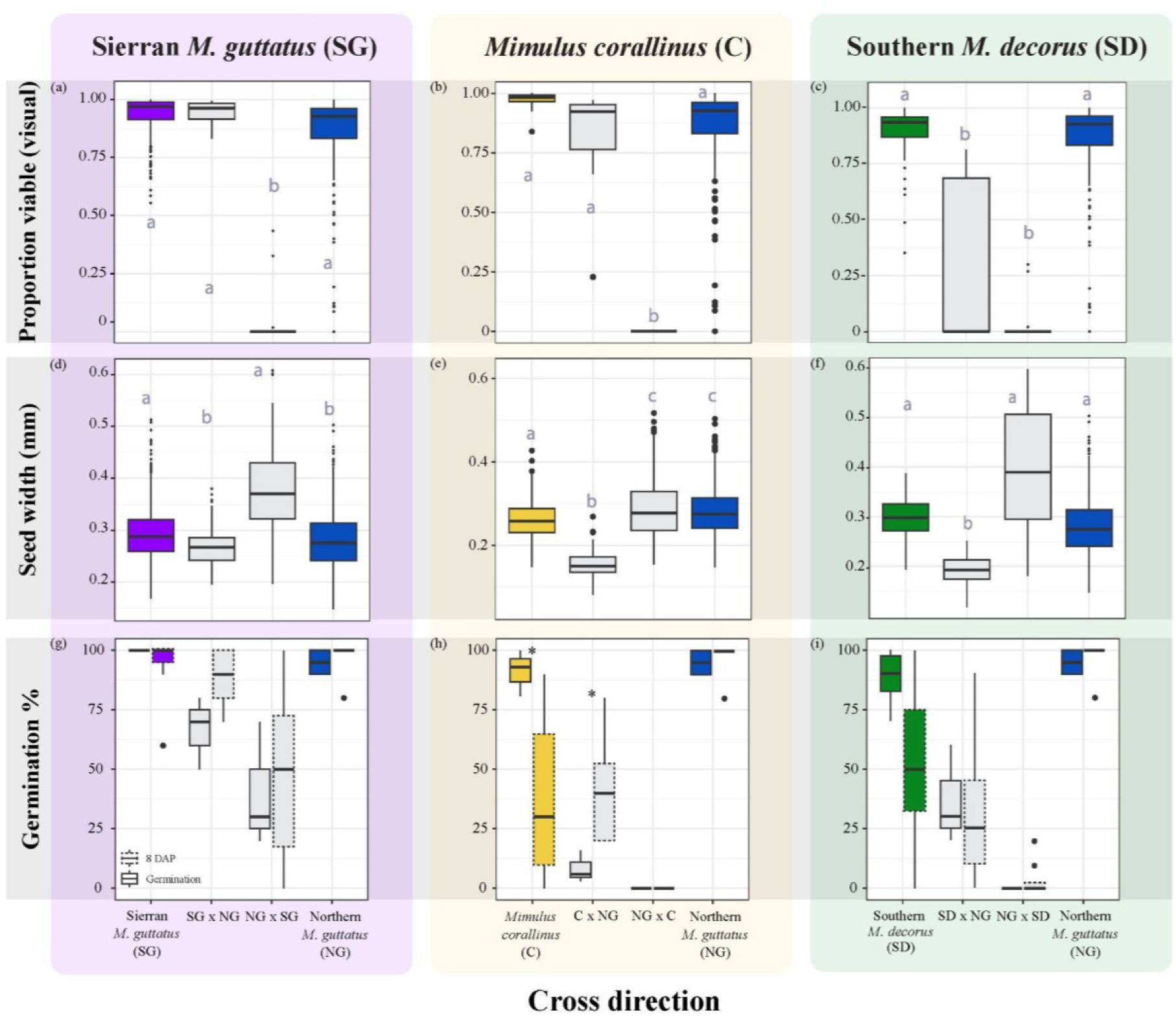
Evidence for parental conflict. Crosses between Sierran *M. guttatus* (SG), *M. corallinus* (C), and Southern *M. decorus* (SD) to Northern *M. guttatus* (SG; columns). **A-C)** The proportion of viable seeds for reciprocal crosses. **D-F)** The seed width for all reciprocal crosses. **G-I)** The result of germination assays (solid outline) and embryo rescued 8 days after pollination (8 DAP; dotted outline) for reciprocal crosses. Groups with significant differences are denoted by letters in plots A-F and by asterisks in plots G-I.

These patterns of hybrid seed inviability mirror those previously described in the *M. guttatus* species complex involving a phylogenetically distinct and allopatric species: Southern *M. decorus* (Fig.1A; Coughlan et al. 2020). Crosses between Southern *M. decorus* and a subset of 9 Northern and Coastal *M. guttatus* lines used in the crossing survey above reveal a broadly consistent pattern, although the pattern here is more symmetric than we have previously reported (Fig.2A-C; Coughlan et al. 2020). In all cases, hybrid seeds are more likely to be inviable when *M. corallinus*, *M. glaucescens,* Sierran *M. guttatus*, or Southern *M. decorus* serve as the sire in crosses with Coastal or Northern *M. guttatus*. Although lines of Coastal and Northern *M. guttatus* did vary from one another in the severity of incompatibility (Fig.1C, Fig.S2), we note that patterns of crossability between any given accession of *M. guttatus* with Sierran *M. guttatus* or Southern *M. decorus* were highly correlated (i.e. lines that exhibit more symmetric patterns of incompatibility with Sierran *M. guttatus* also show more symmetric patterns of incompatibility with Southern *M. decorus*; r^2^ = 0.79, DF = 13, *p* = 0.0005; Fig.S3). We next sought to test whether patterns of repeated hybrid seed inviability supported a role of parental conflict in its evolution.

### Parent-of-origin effects on growth partially support a role of parental conflict

Parental conflict predicts rapid and repeated evolution of hybrid seed inviability through continuous antagonistic selection between maternal- and paternal-interests in resource allocation to offspring (Haig and Westoby 1991). Under parental conflict, hybridization should reveal paternally-acting, resource-acquiring alleles and maternally-acting, resource-repressive alleles. We therefore predict that reciprocal crosses should exhibit parent-of-origin-specific growth phenotypes (Haig and Westoby 1989, 1991; Haig 2000). In line with this prediction, we identified distinct parent-of-origin growth effects in reciprocal F1 seeds in all crosses, wherein seeds sired by *M. corallinus, M. glaucescens,* Sierran *M. guttatus,* or Southern *M. decorus* were larger than the reciprocal cross (Fig. 2D-F; Fig. S4). Given these signatures of paternal excess when *M. corallinus, M. glaucescens,* Sierran *M. guttatus*, or Southern *M. decorus* are the sire, we infer that they have experienced a higher history of conflict than Coastal or Northern *M. guttatus*, in accordance with previous work (Coughlan et al. 2020).

Under parental conflict, disrupted seed development should manifest in the endosperm (Haig and Westoby 1989). We, therefore, predicted that F1 viability may be rescued if embryos are grown on a nutrient-rich media. We performed embryo rescues for a subset of crosses between Northern *M. guttatus* and each of Sierran *M. guttatus, M. corallinus*, and Southern *M. decorus*. While we qualitatively found that germination increased in rescued seeds in 4 out of 6 crosses (Fig.2G-I), only the *M. corallinus* x Northern *M. guttatus* cross was statistically significant (Fig.2H). Despite a lack of statistical significance, we were able to rescue crosses that otherwise never germinate. Moreover, several within-species embryo rescues showed reductions in germination (Fig.2G-I), suggesting that the rescue process itself can be damaging. Therefore, while embryo rescues do not perfectly restore germination, the endosperm likely does play an important role in hybrid seed inviability.

Lastly, parental conflict theory dictates that the outcome of crosses should be predictable based on the histories of conflict, even if phylogenetically distant. In line with this hypothesis, crosses among all high-conflict lineages (i.e. Southern *M. decorus, M. corallinus, M. glaucescens,* and Sierran *M. guttatus*) yielded plump, viable seeds with no parent-of-origin size differences (Fig.3A,B; Fig.S5). Conversely, crosses between high-conflict lineages (i.e. *M. corallinus* and Sierran *M. guttatus*) and a species that has previously been shown to exhibit the lowest history of conflict of all taxa described herein (i.e. Northern *M. decorus;* Coughlan et al. 2020), exhibit reciprocal and nearly complete hybrid seed inviability (Fig.3C,D). Overall, this suggests that high-conflict lineages behave in predictable ways: inviability is most severe between lineages with the most disparate histories of conflict, and viability is recovered between lineages with more similar histories of conflict, despite varying divergence times, in line with parental conflict. However, this prediction may also be true if incompatibility loci have introgressed, a possibility we explore below.

**Figure 3.**
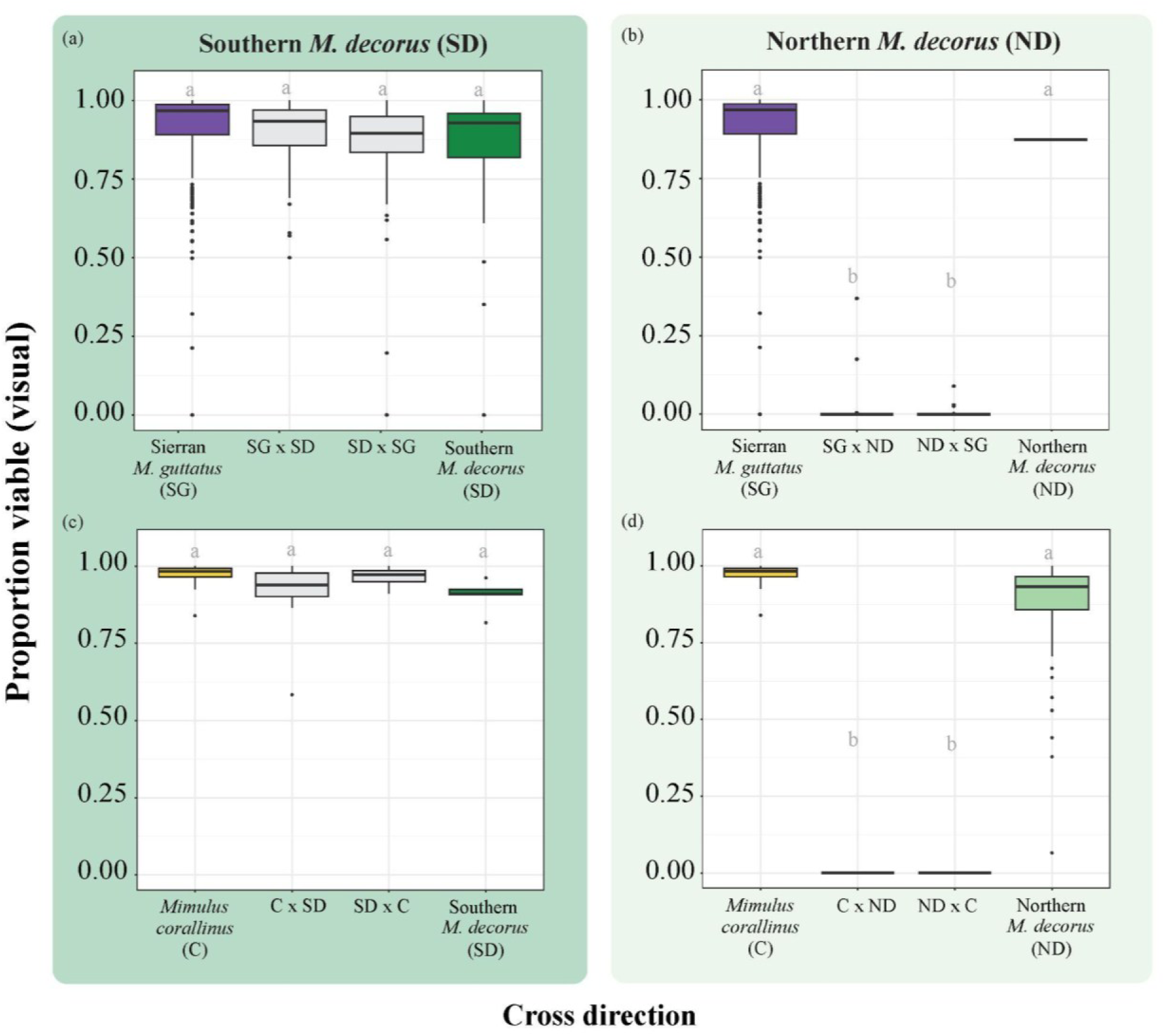
Cross compatibility is predictable based on inferred histories of conflict. High-conflict lineage Southern *M. decorus* is reciprocally crossed to **A)** Sierran *M. guttatus* and **C)** *M. corallinus*, and no hybrid seed inviability is observed. The low-conflict lineage, Northern *M. decorus*, is reciprocally crossed to **B)** Sierran *M. guttatus* and **D)** *M. corallinus*, and nearly complete hybrid seed inviability is observed. Notably, it has been previously shown that the closely related Southern and Northern *M. decorus* have complete hybrid seed inviability when crossed to each other (Coughlan et al. 2020).

### Repeated incidences of hybrid seed inviability have a largely shared genetic basis

We next sought to understand if the shared patterns of cross incompatibility were caused by a shared genetic basis. To test this hypothesis, we used a complementation test, where two lines exhibiting a similar phenotype are hybridized, and the presence or absence of phenotypic segregation in F2 hybrids can reveal whether the genetic basis of the focal trait is shared. Using this logic, we created an F2 population between two lineages that exhibit identical patterns of hybrid seed inviability with Northern *M. guttatus*—*M. corallinus* and Southern *M. decorus* (Fig. 2). We then crossed a single inbred line of Northern *M. guttatus* (IM62) to *M. corallinus*, Southern *M. decorus*, their reciprocal F1s, and reciprocal F2s, with Northern *M. guttatus* acting as the dam in all crosses. If the genetic basis of hybrid seed inviability was largely independent, we would expect that F2s should segregate for the ability to cross with Northern *M. guttatus*, and thus some fraction of F2s would be at least partially compatible with Northern *M. guttatus*. However, we find that out of 761 crosses (>41,000 seeds, 170 unique F2 individuals), we did not recover a single viable hybrid seed, strongly suggesting that the genetic basis of hybrid seed inviability between Northern *M. guttatus* and each of *M. corallinus* and southern *M. decorus* is largely shared (Fig.4A; Fig.S8).

**Figure 4.**
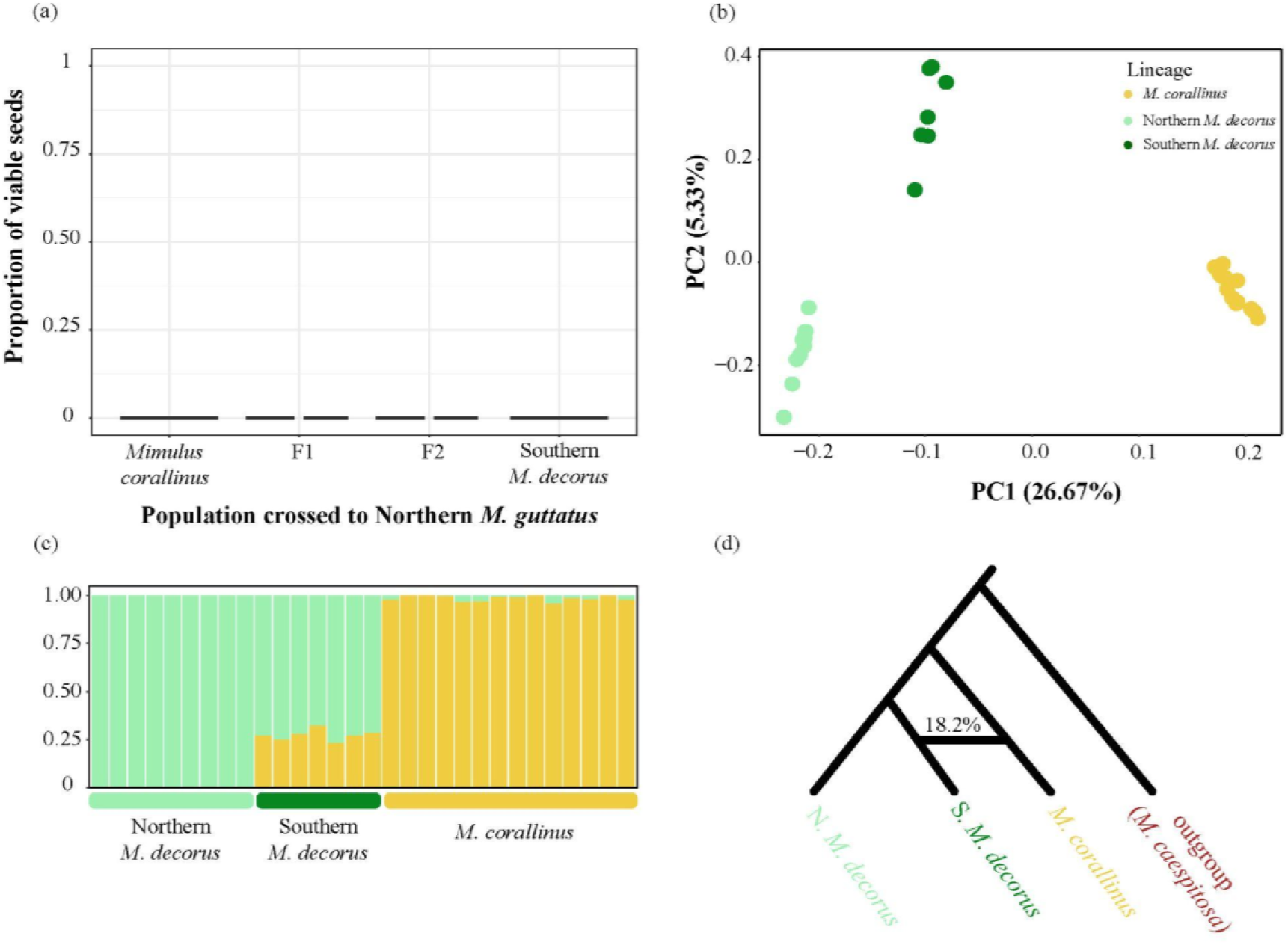
The shared genetic basis of hybrid seed inviability between *M. corallinus* and Southern *M. decorus* may be due to a history of introgression. **A)** A complementation test supports the shared genetic basis of hybrid seed inviability between Northern *M. guttatus* and each of *M. corallinus* and Southern *M. decorus*. **B)** PCA of Northern *M. decorus,* Southern *M. decorus*, and *M. corallinus*. **C)** PCAdmix results for these three lineages. K=2 was the best supported ancestry grouping. **D)** HyDe estimates that 18.2% of the genome of Southern *M. decorus* comes from *M. corallinus*.

### High-conflict lineages exhibit a history of introgression

The shared genetic basis of a trait can be caused by independent mutations in the same or tightly linked gene(s), shared ancestral polymorphisms, or introgression. Given the extensive history of gene flow in this species complex (Brandvain et al. 2014; Twyford and Friedman 2015; Ivey et al. 2023; Mantel and Sweigart 2024; Farnitano et al. 2025), we next tested whether phylogenetically-distinct, high-conflict species may also have a history of introgression. Using four-population tests to examine all combinations of lineages within the *M. guttatus* species complex (with *M. caespitosa* as an outgroup), we find extensive evidence of introgression across the complex, as the minimum value of D is significant in 83% of trios (mean D statistic = 0.059). Moreover, some of these positive tests could be the result of inferred ancient introgression among ancestors of extant lineages (Fig.S6). Given this extensive introgression, disentangling its role in shaping the distribution of hybrid seed inviability is not straightforward, particularly in currently sympatric taxa.

However, there is a particularly strong signal of introgression between two high-conflict lineages— *M. corallinus* and Southern *M. decorus*. These lineages exhibit replicated patterns of hybrid seed inviability with other taxa in the complex (i.e. with both Northern *M. guttatus*, Northern *M. decorus*; Fig.2,3; Coughlan et al. 2020), our complementation test above supports a shared genetic basis between these two lineages for inviability with Northern *M. guttatus*, and they are currently allopatric, potentially allowing for more straightforward tests of introgression. We therefore next focus on investigating the history of ancient introgression between Southern *M. decorus* and *M. corallinus*.

We performed several tests using only the Northern *M. decorus*, Southern *M. decorus,* and *M. corallinus* with *M. caespitosa* as an outgroup. While we sequenced 17 Southern *M. decorus* individuals, we found a high degree of relatedness within populations. We therefore limited most genomic analyses to 6 unrelated individuals. In a PCA, Southern *M. decorus* is intermediate to Northern *M. decorus* and *M. corallinus* along PC1, which explains 26.67% of the genomic variance (Fig.4B). Multiple lines of evidence support a high fraction of *M. corallinus* ancestry segregating within Southern *M. decorus* at a genome-wide level. PCAdmix indicates that there are two ancestries among these three taxa, with Southern *M. decorus* being ∼29% *M. corallinus* and 71% Northern *M. decorus* (Fig.4C). Additionally, we estimated a global admixture proportion of 18.2% *M. corallinus* with HyDe (Fig.4D; Blischak et al. 2018). Using Dsuite, D and f4 also gave broadly consistent estimates of admixture (0.204 and 0.255, respectively; Malinsky et al. 2021). In total, these patterns support a consistent and significant contribution of *M. corallinus* ancestry into Southern *M. decorus*.

We next sought to characterize local patterns of introgression in Southern *M. decorus*. We identified windows exhibiting high *f_dM_*, indicating putative introgression between Southern *M. decorus* and *M. corallinus*. Furthermore, given that the Northern and Southern lineages of *M. decorus* are quite closely related (Coughlan et al. 2020), we should expect relatively low values of F_ST_ between them, except at regions of the genome that have introgressed between *M. corallinus* and Southern *M. decorus.* Such regions should exhibit high F_ST_ between both Northern *M. decorus* and each of *M. corallinus* and Southern *M. decorus*, but low F_ST_ between *M. corallinus* and Southern *M. decorus.* Using these criteria, we find signals of introgression across the genome, typically in small blocks (Fig.5A). These results were consistent with topology weighting (Martin and Van Belleghem 2017), wherein we found support for a phylogeny that grouped *M. corallinus* and Southern *M. decorus* at low levels across the genome, but little support for the phylogeny that grouped *M. corallinus* and Northern *M. decorus*, in line with a scenario of introgression between *M. corallinus* and Southern *M. decorus* rather than incomplete lineage sorting (Fig.S7A). Finally, we used ancestryHMM to infer local ancestry within Southern *M. decorus* (Corbett-Detig and Nielsen 2017); Fig.S7B,C, Fig.S9). Local ancestry across all Southern *M. decorus* is correlated with *f_dM_* (r² = 0.511; p-value < 2.2e-16; Fig.S7D), but reveals high levels of individual ancestry heterozygosity, suggesting that *M. corallinus* alleles are segregating in Southern *M. decorus* populations. Lastly, we estimated the timing of admixture at 4,703 generations (Fig.S10), suggesting that introgression is not ongoing.

**Figure 5.**
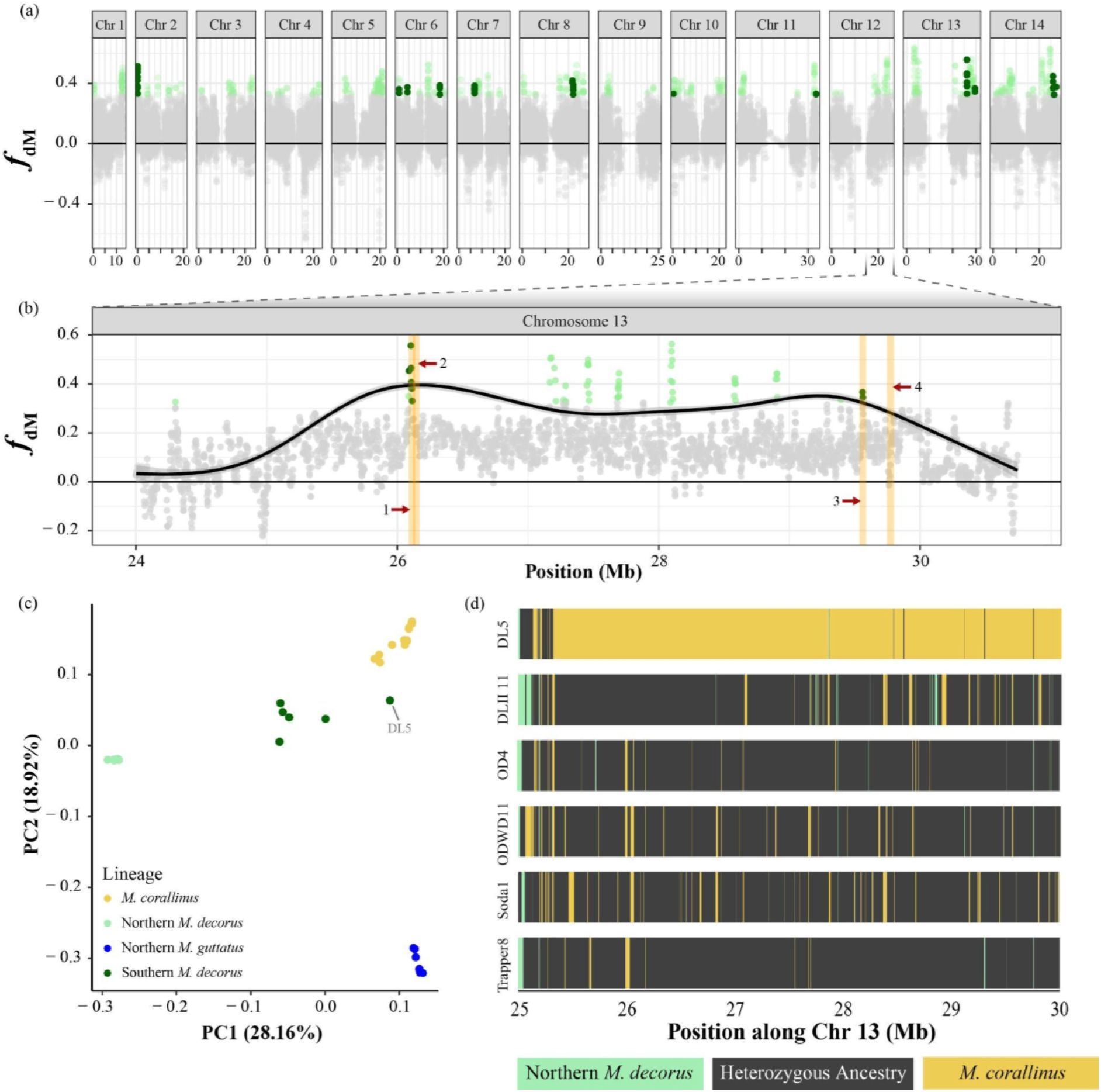
The Southern *M. decorus* genome is a genomic mosaic of Northern *M. decorus* and *M. corallinus*. **A)** Distribution of *f*_dM_ values measured in 250 SNP windows. Light green dots are in the top 1% of *f*_dM_ values. Dark green dots represent the windows that also overlap with the top 1% of windows with low F_ST_ between *M. corallinus* and Southern *M. decorus* and high F_ST_ between northern and southern *M. decorus*. **B)** The putative inversion on chromosome 13. The black line represents the smoothed average of F_ST_ between *M. corallinus* and Southern *M. decorus* minus F_ST_ between Northern and Southern *M. decorus*. Orange blocks indicate the location of genes with known roles in imprinting or endosperm expression: (1) *Napin*, (2) *MSI* homolog, (3) *DOG1*, and (4) *FIE* homolog. **C)** A PCA of SNPs within the inversion region places most Southern *M. decorus* samples intermediate to Northern *M. decorus* and *M. corallinus*, except for DL5, which falls closer to *M. corallinus*. **D)** Each unrelated Southern *M. decorus* sample is colored by the most likely ancestry at each position, determined by AncestryHMM. DL5 is largely homozygous for *M. corallinus* ancestry, while the remaining unrelated samples are largely heterozygous.

Although we find many small blocks of introgression in the genomes of Southern *M. decorus*, one striking signal occurs at the distal end of chromosome 13 (Fig.5B). This region exhibits a high *f_dM_*, as well as a high difference in F_ST_ when comparing Southern and Northern *M. decorus* to *M. corallinus*. This region harbors a chromosomal inversion known to be distinct between Northern *M. decorus* and *M. guttatus* (Lovell et al. 2025; Joint Genome Institute, pers. comm.). Intriguingly, this inversion has previously been implicated in hybrid seed lethality between *M. guttatus* and *M. tilingii* (Garner et al. 2016), and preliminary mapping data supports its role in hybrid seed inviability between Northern *M. decorus* and *M. guttatus* (Coughlan unpublished data). Excitingly, this inversion contains multiple genes that are potential targets of parental conflict that are also introgression outliers (i.e. are in the top 1% for *f_dM_* and F_ST_ difference (Table S2)). Two outlier windows contain genes known to be expressed in the *M. guttatus* endosperm (Tucci et al. 2024), although it is currently unknown whether either of these genes are imprinted in *Mimulus*. One top window contains a homolog of *Napin*, a storage protein found in maturing seeds (Crouch et al. 1983), while the other contains the *Mimulus* homolog of *DOG1*, which plays an important role in seed dormancy (Bentsink et al. 2006). There are also two genes that are known to be imprinted in *Arabidopsis*. The first gene is an *MSI* homolog, located close to *Napin* in one of the top windows. Just downstream of the second top window is *FIE*, a gene that is both a master regulator of imprinting in *Arabidopsis* endosperm and is itself imprinted (Luo et al. 1999, 2000; Köhler et al. 2003, 2021; Batista and Köhler 2020).

We hypothesized that this inversion has introgressed from *M. corallinus* into Southern *M. decorus*, and may play an important role in the nearly complete hybrid seed inviability between the two sister lineages of *M. decorus*. However, we note that the role of this inversion is not straightforward and cannot explain all patterns of hybrid seed inviability that we describe in the species complex as *M. corallinus* and many *M. guttatus* lineages share the same orientation, despite also exhibiting hybrid seed inviability (Lovell et al. 2025). Additionally, we find that the inversion is heterozygous in 15 out of 17 Southern *M. decorus* samples (5 out of 6 unrelated samples). The observation that this inversion contributes to hybrid seed inviability in other *Mimulus* crosses but is found largely in heterozygous state may seem perplexing, as segregation of the inversion in the offspring of Southern *M. decorus* should result in segregating hybrid seed inviability– a pattern that we do not see. However, the observed patterns of hybrid seed inviability may be explained if the genetic basis of hybrid seed inviability between the two lineages of *M. decorus* is sufficiently redundant (i.e. composed of several large affect alleles with redundant effects). Although we find strong evidence of introgression of this inversion, it contains potential targets of parental conflict, and is known to contribute to hybrid seed inviability in other crosses, further genetic mapping will be required to explicitly link this chromosomal inversion with hybrid seed inviability between the two lineages of *M. decorus*.

## Discussion

We show that hybrid seed inviability is widespread and has evolved rapidly and repeatedly within the *Mimulus guttatus* species complex. Hybrid seed inviability segregates among the widespread *M. guttatus*, as well as several other species in the group, yet patterns of hybrid seed inviability are more strongly associated with geography, rather than phylogeny. Crossing patterns within this group also conform to predictions of parental conflict theory: crosses exhibit canonical parent-of-origin effects on growth, incompatibility is at least partially linked with the nutritive endosperm, and patterns of crossing are more predictive based on the inferred history of conflict that lineages have experienced, rather than phylogenetic relatedness. Lastly, two instances of hybrid seed inviability appear to have a shared genetic basis, and these two taxa also exhibit a substantial history of introgression, including at several candidate genes. Although more research is needed to explicitly identify genes involved in hybrid seed inviability and assess their histories of introgression, our results highlight the potential for intragenomic conflicts to drive the rapid evolution of incompatibility alleles, and for introgression to underlie the invasion of incompatibility alleles into new species. We posit that introgression of selfish alleles could then cause cascading reproductive isolation between populations of what was once the same species.

### Parental conflict as a driver of hybrid seed inviability

The data we have presented here is consistent with the parental conflict theory. This theory posits an ongoing arms race between paternally-acting, resource-acquiring alleles and maternally-acting, resource-repressive alleles within populations. When populations with different histories of conflict hybridize, incompatibilities between these alleles are revealed. A hallmark phenotype of parental conflict is parent-of-origin growth defects in F1 hybrids, which is a consistent observation in all crosses. Furthermore, parental conflict theory predicts that incompatibilities should manifest in endosperm. Therefore, it should be possible to “rescue” embryos by providing them with an external source of nutrition. Our embryo rescue results are less conclusive than the patterns in seed size, but they trend in the expected direction, and we were able to rescue hybrid embryos in crosses that typically do not germinate. Moreover, the procedure itself is delicate, as exemplified in our control embryo rescues (which in some cases are significantly worse than the regular germination assay; Fig. S2), suggesting that our partial hybrid rescue may also be influenced by the rescue procedure itself. Lastly, we find that the inferred histories of conflict are highly predictive of the outcome of future crosses. Despite being more distantly related, all Sierran lineages were entirely compatible with one another, as well as with high-conflict Southern *M. decorus*. By contrast, all taxa were completely incompatible with the lower-conflict lineage, Northern *M. decorus*.

The introgressed chromosome 13 inversion is also connected to these predictions. Several genes located in or near our top *f*_dM_ peaks within this region are expressed in the *Mimulus* endosperm (e.g., *DOG1* and *Napin;* Tucci et al. 2024) or imprinted in *Arabidopsis* and contribute to parent-of-origin effects when disrupted in that species (e.g., *FIE* and *MSI*; Luo et al. 1999; Köhler et al. 2003). In addition to being imprinted themselves, both *FIE* and *MSI* function as part of the polycomb repressive complex, which establishes the histone modifications needed for imprinting and proper endosperm development (Mozgova et al. 2015; Batista and Köhler 2020). Moreover, this region has been implicated in hybrid seed inviability in at least two crosses (Coastal *M. guttatus* x *M. tilingii* and Northern *M. guttatus* x Northern *M. decorus*; Garner et al. 2016; Coughlan unpublished data). Although some of our observations give rise to more questions (i.e. excess levels of heterozygosity within the inversion), we hypothesize that introgression of this inversion from *M. corallinus* into what is now Southern *M. decorus* led to cascading reproductive isolation with what is now Northern *M. decorus*. Nonetheless, explicit tests of its role in generating hybrid seed inviability between Southern *M. decorus* and *M. guttatus* will require additional genetic mapping.

Our work adds to a growing body of evidence for parental conflict across diverse taxa (Long 2005; Schrader and Travis 2008; Chuong et al. 2010; Babak et al. 2015; Cailleau et al. 2018; Raunsgard et al. 2018; Coughlan et al. 2020; Sandstedt and Sweigart 2022; Petrén et al. 2023; Butel et al. 2024). Our results of widespread, asymmetric hybrid seed inviability are not only consistent with other studies in diverse angiosperms, but also mirror the parent-of-origin phenotypes observed in mammals, where the placenta performs an analogous function to the endosperm (Gutiérrez-Marcos et al. 2004; Feil and Berger 2007; Vrana 2007; Briscoe Runquist et al. 2014; Rebernig et al. 2015; Oneal et al. 2016; Lafon-Placette et al. 2017, 2018; Roth et al. 2018, 2019; Arévalo and Campbell 2020; Coughlan et al. 2020; Brekke et al. 2021; Florez-Rueda et al. 2021; İltaş et al. 2021; Sandstedt and Sweigart 2022; Soliman and Coughlan 2024). These parallels suggest that parental conflict could be a common driver of early-onset hybrid embryonic death, especially in cases where isolation is rapidly evolving and/or actively segregating, as is the case in the *M. guttatus* species complex.

### A polymorphic incompatibility sorted by geography

One striking finding from our work is that hybrid seed inviability segregates within the widespread *M. guttatus*. Polymorphic incompatibilities are common in nature (Corbett-Detig et al. 2013; Sicard et al. 2015; Larson et al. 2018; Zuellig and Sweigart 2018; Calvo-Baltanás et al. 2021), and may represent incompatibilities en route to fixation (or loss; Cutter 2012). Yet, it is often unclear how such alleles would avoid removal from the population, especially when rare (Bank et al. 2012; Lindtke and Buerkle 2015; Xiong and Mallet 2022). If the alleles themselves are selfish, such as targets of parental conflict, the selective advantage conferred by these alleles may aid in overcoming their deleterious epistatic effects (Werren 2011; Meiklejohn et al. 2018; Wang et al. 2024). In the case of other selfish elements, such as meiotic drivers or *Wolbachia*, much theory and empirical work has demonstrated the ability of such selfish loci to invade and spread, even in the face of deleterious effects to the host (Hoffmann et al. 1990; Raychoudhury et al. 2009; Werren 2011; Meiklejohn et al. 2018; Turelli et al. 2018), but an equivalent body of work is lacking in the case of alleles evolving via parental conflict.

Additionally, these patterns of hybrid seed inviability do not align with inferred phylogenetic relationships presented here or alternative topologies presented elsewhere (Whitener et al. 2024). For example, all Sierran *M. guttatus* have a similar crossing pattern and are compatible with one another, yet they are not a monophyletic group (Fig.1A). Instead, we identify the Sierra mountains and foothills as a unique group that exhibits cross incompatibility with other geographic regions of *M. guttatus*, but broad compatibility with themselves. Although the *M. guttatus* species complex inhabits a broad range from Alaska to Mexico and from Pacific coastline through to the Rocky mountains (Vickery 1974, 1978), the Sierras represent a unique geographic location where two deeply diverged clades within this complex meet (i.e. Northern and Southern *M. guttatus;* Twyford and Friedman 2015), and is home to several endemic species (Ferris et al. 2014, 2017; Twyford and Friedman 2015; Coughlan et al. 2021; Ivey et al. 2023). Although more work on the phylogeography of this group will be needed to further assess the history of the Sierras, one compelling hypothesis is that the Sierras may have contained unique refugia during the last glacial maximum, allowing for population divergence and the accumulation of incompatibility alleles. After glacial retreat and range expansion, such incompatibility alleles may have spread to nearby Northern *M. guttatus* or *M. glaucescens* populations in the Sierra foothills. In line with this hypothesis, we find excessive allele sharing between the Northern and Southern clades of *M. guttatus* in the Sierra mountains and foothills (Fig.S6). Cases of complex population structure and high diversity driven by microrefugia within the Sierras are known in other species (Weng et al. 2021). Alternatively, these incompatibility alleles may predate the split of the Northern and Southern clades, and may have independently fixed in the Sierra mountains and foothills. Given that the entire complex exhibits significant allele sharing some of which is caused by introgression (Brandvain et al. 2014; Twyford and Friedman 2015; Ivey et al. 2023; Mantel and Sweigart 2024), distinguishing patterns of incomplete lineage sorting from introgression within *M. guttatus* is challenging and will require a much deeper understanding of the relationship among populations in the Sierras and beyond (though see: Twyford and Friedman 2015; Twyford et al. 2020). Despite this, one clear incidence of introgression is that of *M. corallinus* into the now allopatric Southern *M. decorus*, the consequences of which we explore below.

### Introgression as a source of incompatibilities

Our work supports a shared genetic basis of hybrid seed inviability and a history of introgression between *M. corallinus* and Southern *M. decorus*. Given the low divergence between Southern *M. decorus* and its sister species, Northern *M. decorus*, we infer that this introgression may have led to cascading reproductive isolation between the two *M. decorus* lineages.

Yet, many questions remain about this case of introgression. First, the current geographic distance between *M. decorus* and *M. corallinus* suggests that introgression is not ongoing, but it is unclear how and where this introgression originally occurred. The distribution of the *M. guttatus* species complex is likely a result of post-glaciation recolonization, particularly in the Sierras (Twyford and Friedman 2015), which may have facilitated hybridization. In line with this idea, our estimate of the timing of admixture corresponds to a period of geological dynamism in the southern Cascades, wherein many of the current Cascade Lakes were formed (4,703 generations; 4-14kya, assuming a 1-3 year generation time; Johnson 1985). It is, of course, also possible that the source of introgression was a close relative of *M. corallinus* that was not sampled or is no longer extant. Indeed, regions that we identified as fixed for *M. corallinus* ancestry within southern *M. decorus* are not identical (D*_xy_* = 0.017; F_ST_ = 0.007). Still, even with the high diversity of this species complex (Lovell et al. 2025), our data support *M. corallinus* as the most likely source of the introgression (Fig. S6).

Introgression may be an underappreciated contributor to reproductive isolation. Although introgression is often viewed as a homogenizing force (Owens and Samuk 2020), it can also generate novel biodiversity (Otto and Whitton 2000; Lamichhaney et al. 2018; Long and Rieseberg 2024; Rosser et al. 2024). However, in between these extremes, there are opportunities for lower levels of introgression to move incompatibility alleles between populations (Widen et al. 2023; Aguillon et al. 2025). In the case of Northern and Southern *M. decorus* presented in this study, it is possible that introgressed alleles from *M. corallinus* provided an advantage in parental competition to isolated populations of *M. decorus*, leading to the origin of the Southern lineage. We have previously shown that seeds developing alongside hybrids from low-conflict fathers are larger than seeds growing beside full siblings (Coughlan 2023a), suggesting that at the level of individual alleles, selfish solicitation of maternal resources confers a direct benefit to the individual seed (and a direct cost to sibling seeds). Yet, how these small increases in seed size translate into survival and fitness in the wild remains unknown. Few incidences of introgressed incompatibilities have been reported, but given the ubiquity of introgression across eukaryotes (Taylor and Larson 2019; Dagilis et al. 2022), they may be more prevalent than previously thought, as recently exemplified in Swordtail fishes (Aguillon et al. 2025), as well as a potential example of chloroplast capture in a more distantly related group of monkeyflowers (Nelson et al. 2021).

## Conclusion

Overall, we find that reproductive isolation is segregating in the *Mimulus guttatus* species complex, following a geographic rather than phylogenetic pattern. In at least one instance, we have provided evidence that introgression may have moved incompatibility alleles between lineages, generating reproductive isolation between sister lineages. Our study supports the idea that parental conflict and introgression of selfish elements can contribute to the rapid, repeated, and strong evolution of intrinsic postzygotic reproductive isolation.

## Methods

### Crossing surveys

To assess hybrid seed inviability within the widespread *M. guttatus*, we grew 3-5 individuals from 1 inbred line/maternal family for each of 23 populations of *M. guttatus,* which span a large part of the species’ geographic range (Fig. 1B; Table S1). Seeds were cold stratified at 4°C for 5 days before being transferred to long days in the University of North Carolina greenhouses (16hrs light, 22°C). Upon flowering, we crossed all accessions to each other in a nearly full diallel design, totalling 426 unique cross directions (out of a possible 506), with a mean of 4 replicate crosses per combination (range: 1-12, totalling 1,742 crosses). For all crosses, the buds were emasculated 1-2 days before opening to prevent autogamous self-fertilization, and pollen from a freshly opened flower of the sire was applied to a receptive stigma of the dam.

Based on the results of these crosses, we performed follow-up crossing surveys that incorporated other species within the complex: *Mimulus decorus*, *M. corallinus*, and *M. glaucescens*. Previously, we found that *M. decorus* was two genetically unique lineages, Northern and Southern, that vary in their ability to cross with *M. guttatus*, putatively via differences in the strength of parental conflict within each lineage (Coughlan et al. 2020). Therefore, we crossed 12 accessions of *M. guttatus* (which spanned the observed variation in crossing pattern in the initial survey) to 4 populations of Southern *M. decorus* (high-conflict lineage; mean of 4 replicate crosses/direction, 730 crosses total), and 3 populations of Northern *M. decorus* (low-conflict lineage; mean of 3 replicate crosses/direction, 256 crosses total). For *M. glaucescens,* we crossed individuals from 2 populations to 21 *M. guttatus* populations used in the original survey, as well as 2 populations of Northern *M. decorus* and 4 populations of Southern *M. decorus* (mean of 4 replicate crosses, 458 crosses total). For *M. corallinus*, we crossed 3 accessions of *M. corallinus* to one accession of each Northern *M. guttatus,* Sierran *M. guttatus*, Northern *M. decorus*, and Southern *M. decorus* (3 replicate crosses/direction, 95 crosses total).

### Estimation of seed viability, germination, and size

We estimated seed viability in two ways. First, we counted all seeds formed per fruit and characterized viability based on morphology (e.g., shriveled/flat corresponding to inviable, or round corresponding to viable). To confirm germination proclivity, we performed a germination assay where 10 seeds from each unique cross combination were plated on 0.6% agar, cold stratified for 1 week, then placed in a growth chamber under long-day conditions (18hrs light, 21/18°C day/night) at Yale University. We seeded 3 replicated plates per unique cross. Plates were monitored weekly for 4 weeks for the emergence of a radicle, which was deemed a germinant. These two measurements of viability are highly correlated (r^2^ = 0.81, DF = 532, *p* < 0.0001), but offer different insights; while one characterizes abnormal phenotypes, the other directly assesses death. To determine seed size, we photographed seeds on white paper with a Nikon D3500 DSLR digital camera and a size standard. We measured the width of 25 seeds for each fruit manually using ImageJ (Schneider et al. 2012). To assess cross compatibility and parent-of-origin differences in seed size, we ran a series of linear mixed models using the *lme4* library in R (Bates et al. 2015), where either the proportion of viable seeds or seed width was the response variable, cross type was a fixed effect, and the identity of the maternal and paternal populations were each random effects. In both sets of analyses, cross type contained four levels (within-lineage for each parent and each reciprocal F1). For each model, we then assessed differences among cross types with a type III Anova using the *car* package and identified significant comparisons using contrasts from the *emmeans* package in R (Fox and Weisberg 2019; Lenth 2025).

### Embryo Rescue

We further wanted to determine if hybrid seed inviability was caused by defects in the endosperm rather than the embryo itself. To test this, we performed an embryo rescue experiment where early developing seeds were grown on Murashige and Skoog media supplemented with 4% sucrose to functionally compensate for the endosperm by providing external nutrition needed for growth. Under long days at the Yale University greenhouses, we grew four individuals from a single inbred line representing each of the four species used in this assay: Northern *M. guttatus* (IM62), Sierran *M. guttatus* (CCC9), *M. corallinus* (SILF), and Southern *M. decorus* (OD11). Given that all incidences of hybrid seed inviability reported here involve Northern *M. guttatus*, we crossed Northern *M. guttatus* reciprocally with the other three species and included intraspecific crosses as controls. For each cross direction, we performed three replicate crosses and dissected the ovaries eight days after pollination (DAP). Ten seeds per cross were placed on Petri dishes with media, sealed with 3M micropore tape, then incubated in growth chambers under long-day conditions (18 hrs light, 22/20°C day/night). Seeds were scored for germination weekly for four weeks. For each cross direction, we tested for significant difference between the germination assay and 8 DAP embryo rescue using unpaired non-parametric Wilcoxon rank-sum test in R.

### Complementation test

In this study, we report multiple incidences of hybrid seed inviability and shared patterns of incompatibility across phylogenetically and geographically distinct taxa. Given that we find the repeated occurrence of high-conflict lineages within the *M. guttatus* complex, we next sought to understand if the genetic basis of the incompatibility was shared among incidences. To do so, we performed a complementation test between two geographically distinct taxa that showed similar patterns of hybrid seed inviability when crossed with Northern *M. guttatus*-Southern *M. decorus* and *M. corallinus*. We first crossed these two species reciprocally, then self-fertilized one reciprocal F1 from each cross direction to create reciprocal F2 populations. We predict that if the genetic basis of hybrid seed inviability with Northern *M. guttatus* was largely shared between *M. corallinus* and Southern *M. decorus*, all F2s should produce inviable seeds when acting as the sire in crosses with Northern *M. guttatus*, just as we see for both *M. corallinus* and Southern *M. decorus.* By contrast, if the genetic basis was independent, then F2s should segregate for their crossability with Northern *M. guttatus*, resulting in some fraction of F2s that produced viable hybrid seeds. To test these predictions, we planted 5 replicates of each parent, along with 15 F1s from each cross direction, and roughly 100 F2s from each cross direction. After omitting individuals that did not flower, we were left with 3 Southern *M. decorus*, 4 *M. corallinus*, 15 Southern *M. decorus* x *M. corallinus* F1s, 12 *M. corallinus* x Southern *M. decorus* F1s, and 170 F2s (83 with Southern *M. decorus* as the grand-dam and 87 with *M. corallinus* as the grand-dam). We crossed each experimental plant with an inbred line of Northern *M. guttatus* (IM62), with IM62 serving as the dam and each experimental plant as the sire for all crosses. We performed an average of 3 replicate crosses per plant (range: 1-10), totalling 761 crosses. Based on our previous results, hybrid seed inviability should manifest as large, flat, disc-like seeds. We assessed the percentage of viable seed per fruit.

Additionally, we performed a simplified power analysis to assess under what genetic conditions we would be able to detect segregating phenotypes among our F2s. For all models, we assumed *x* unlinked loci with equivalent and additive effect sizes, and assessed the expected fraction of F2 individuals that would exhibit complete inviability with *M. guttatus*. Here, *x* varied from 1 to 10 loci, and we simulated four levels of effect size: each locus having a 100% lethal effect, 50% lethality, 25% lethality, or 10% lethality. We then performed a Fisher’s Exact test between our observed counts of individuals that produced no viable seeds (versus those that produced at least some viable seeds) with what would be expected under each set of genetic conditions. We find that, except under the most extreme parameters (i.e. 6 or more loci, each of which exhibits 100% lethality or 9 or more loci each of which exhibits 50% lethality; Fig. S8), we have the power to identify a lack of segregation among F2s. We note that these are quite conservative estimates, as other studies have found that hybrid seed inviability typically involves a smaller number of paternally acting alleles (i.e. 3-6), each of which reduces viability by 20-50% (Rebernig et al. 2015; Garner et al. 2016; Lafon-Placette et al. 2017; Couglan unpublished data).

### Genomic Analyses

In order to contextualize the patterns observed above with the underlying genetic relationships between lineages, we used whole genome sequences from the lines used in our experiments, combined with numerous new and publicly available sequences. We sequenced 99 new accessions and combined this with 73 publicly available accessions (Table S1). For new sequences, we extracted DNA from bud and leaf tissue using the ThermoFisher GeneJET Plant Genomic DNA kit. Illumina libraries were prepped and sequenced using NovaSeq6000 by the Yale Center for Genome Analysis. For all accessions, we cleaned reads using fastp v0.23.2 (using option “-q 20”; Chen et al. 2018) and aligned to the IM62v3 reference genome (Lovell et al. 2025) using the BWA mem algorithm v0.7.17 (Li and Durbin 2009). We removed duplicates using Picard v2.25.6 (options “VALIDATION_STRINGENCY=LENIENT”; http://broadinstitute.github.io/picard). We called variants using bcftools mpileup and call (v1.16; Li 2011; Danecek et al. 2021), and filtered for quality and depth (-e “FMT/DP<4 | FMT/GQ<20 | QUAL<40”) using bcftools v1.16.

We estimated relatedness among samples using king (v 2.3.2; Manichaikul et al. 2010). Next, we constructed a phylogeny for all lineages using IQTree2 (Minh et al. 2020). We removed individuals with less than 3X coverage, as well as those suspected to be early generation hybrids, leaving 158 sequences. We used only biallelic sites and sites with a minimum minor allele frequency of 0.05. We selected the best substitution model, TVM+F+ASC+R6, using ModelFinder (Kalyaanamoorthy et al. 2017), and then we used that model to infer a consensus tree using 1000 bootstraps (Fig. 1A).

In order to broadly assess diversity, divergence, and introgression among these lineages, we calculated D and F4 statistics using Dsuite (Malinsky et al. 2021). We split samples into their respective lineages by combining field identification with the inferred relationships from the phylogeny, as well as PCAdmix run using all samples. We ran all combinations of *M. guttatus* species complex lineages with Dtrios using a more distantly related and allopatric species-*M. caespitosa-* as an outgroup. We compared the arrangement of the trios that minimized the value of D to the arrangement predicted by the phylogeny generated above. We also ran Dtrios constrained by the phylogeny, and used these results to obtain f-branch values (Malinsky et al. 2021).

We further compared the genomes of the two high-conflict lineages for which we had performed the complementation test (*M. corallinus* and Southern *M. decorus)*, as well as the low-conflict sister species of Southern *M. decorus*: Northern *M. decorus*. We used PCANGSD (Meisner and Albrechtsen 2018) to perform PCA and PCAdmix for global ancestry grouping. Based on these initial findings, we found a striking pattern of ancestry sharing between the two high-conflict lineages. To investigate this shared ancestry further, we constructed explicit tests of introgression using Dsuite and calculated admixture proportion for southern *M. decorus* using HyDe (Blischak et al. 2018), using standard settings with *M. caespitosa* as an outgroup.

In order to investigate patterns of introgression along the genome, we used Dinvestigate to calculate *f*_dM_ in sliding windows of 250 SNPs with a 50 SNP step. We selected fourfold degenerate sites using degenotate (Mirchandani et al. 2024), and used these to calculate F_ST_ with *Pixy* (Korunes and Samuk 2021) using the same sliding windows in which *f*_dM_ was calculated. We further investigated regions of the genome that appear introgressed by estimating topology weighting along the genome using TWISST (Martin and Van Belleghem 2017). TWISST input files were made using the genomics_general scripts available from Simon Martin (https://github.com/simonhmartin/genomics_general), as recommended by the TWISST manual, again using *M. caespitosa* as an outgroup.

To understand the individual-level patterns of introgression, we inferred local ancestry within our Southern *M. decorus* genomes using ancestryHMM (Corbett-Detig and Nielsen 2017). We used all *M. corallinus* and Northern *M. decorus* individuals to calculate population allele frequencies for the parental populations using custom scripts. To assign genetic distances between loci, we used a linkage map generated from crosses between Northern *M. decorus* and *M. guttatus* (Schwarz, unpublished); for sites in between SNPs on the linkage map, we assumed a linear increase in genetic position with physical position. We used a starting ancestry proportion of 20% *M. corallinus* ancestry and allowed the timing of that ancestry to be inferred by the program. We compared three different sets of SNPs: 1) all sites that had variation in the parental populations, 2) fixed differences between the parents, and 3) sites with an allele frequency difference between the parents greater than 50%. We only included SNPs that had no missing data in the parental samples (Fig.S9). We focus on the results from using all variable sites as these analyses yielded the most consistent results with other genomic analyses.

Finally, our *M. corallinus* samples displayed significant population structure, with four individuals forming a distinct group in the phylogeny and PCA. Therefore, we repeated all of our analyses with and without those individuals. There was not a substantial difference in the results, so we present only the results using all *M. corallinus* individuals here.

## Author Contributions

All authors contributed to the design and conceptual framing of this work, as well as data collection and data analysis. M.E.F. led the writing of the manuscript with significant contributions from all authors. All authors approved the final version of this manuscript before submission.

## Acknowledgements

We are grateful to Christopher Bolick and Nathan Guzzo for plant care and assistance in the Yale Science Building greenhouses. The Coughlan lab provided helpful feedback on earlier drafts of this work. We also thank Patricia Vallejo Joseph and Quinn Evans for helping with seed counting and viability assessment. This work was supported by an NIH grant to J.M.C. (NIH R35GM150907), an NSF Postdoctoral Research Fellowship in Biology to M.E.F. (2305853), and an NIH grant to Daniel R. Matute that supported J.M.C. (NIH R35GM148244). P.F.S. and H.K.S. were supported by the Yale Graduate School. The Yale Center for Genome Analysis, which provided sequencing, is supported by the NIH NIGMS 1S10OD030363-01A1.

