## Supplemental Tables and Figures for "Introgression and Parental Conflict Shape Repeated Occurrences of Postzygotic Isolation"

**Table S1. Accessions used for crossings and genomic analyses.**

| Accession | Lineage | SRA Number | Used in Crosses? | Used in Genomics? |
|---|---|---|---|---|
| **AHQT** | Northern *M. guttatus* | SRR486613 | No | Yes |
| **ASPEN1** | *M. corallinus* | SRR35572364 | No | Yes |
| **BCB2** | Coastal *M. guttatus* | SRR35572363 | Yes | Yes |
| **BID2nas** | *M. nasutus* | SRR35572352 | No | Yes |
| **BID6nas** | *M. nasutus* | SRR35572341 | No | Yes |
| **BOG10** | Northern *M. guttatus* | SRX030570 | Yes | Yes |
| **BOG4** | Northern *M. guttatus* | NA | Yes | No |
| **BR4** | Northern *M. decorus* | SRR35572330 | No | Yes |
| **BR5** | Northern *M. decorus* | SRR23709133 | No | Yes |
| **CACG6** | Northern *M. guttatus* | SRR1259270 | No | Yes |
| **CACN9** | *M. nasutus* | SRR1259271 | No | Yes |
| **CCC9** | Sierran *M. guttatus* | SRR35572319 | Yes | Yes |
| **CCR13** | Sierran *M. guttatus* | SRR35572308 | No | Yes |
| **CCR18** | Northern *M. guttatus* | SRR35572297 | No | Yes |
| **CCR4** | Sierran *M. guttatus* | SRR35572286 | Yes | Yes |
| **CCR8** | Northern *M. guttatus* | SRR35572275 | No | Yes |
| **CF3r12** | *M. corallinus* | SRR35572362 | No | Yes |
| **CH30glau** | Northern *M. guttatus* | SRR35572361 | No | Yes |
| **CHM10** | Northern *M. guttatus* | SRR35572360 | No | Yes |
| **CHM4** | Northern *M. guttatus* | SRR35572359 | No | Yes |
| **CHM7** | Northern *M. guttatus* | SRR35572358 | No | Yes |
| **CHR11Gu** | Northern *M. guttatus* | SRR35572357 | No | Yes |
| **CHR18nas** | *M. nasutus* | SRR35572356 | No | Yes |
| **CHR41nas** | *M. nasutus* | SRR35572355 | No | Yes |
| **CHR8** | Northern *M. guttatus* | SRR35572354 | No | Yes |
| **Cohass14** | *M. nasutus* | SRR35572353 | No | Yes |
| **Cohass1n** | *M. nasutus* | SRR35572351 | No | Yes |
| **ConCrk15** | *M. nasutus* | SRR35572350 | No | Yes |
| **CRES3** | Northern *M. guttatus* | SRR35572349 | No | Yes |
| **CSS4** | Southern *M. guttatus* | SRR9674909 | No | Yes |
| **Cultus1** | Northern *M. guttatus* | SRR35572348 | No | Yes |
| **CVP9** | Southern *M. guttatus* | SRR486607 | No | Yes |
| **CWF30** | Northern *M. guttatus* | SRR35572347 | No | Yes |
| **DEB9** | Northern *M. guttatus* | SRR35572346 | No | Yes |
| **DES6** | Northern *M. guttatus* | SRR35572345 | No | Yes |
| **Devil27** | Northern *M. guttatus* | SRR35572344 | No | Yes |
| **DINKEY2** | Southern *M. guttatus* | SRR35572343 | No | Yes |
| **DL1** | Southern *M. decorus* | SRR10194633 | No | Yes |
| **DL5** | Southern *M. decorus* | SRR35572342 | Yes | Yes |
| **DLII_11** | Southern *M. decorus* | SRR35572340 | No | Yes |

| | | | | |
|---|---|---|---|---|
| **DM1** | *M. corallinus* | SRR35572339 | No | Yes |
| **DPRN** | *M. nasutus* | SRR1298375 | No | Yes |
| **EAG** | *M. corallinus* | SRR13618753 | Yes | Yes |
| **EAM** | *M. corallinus* | SRR35572338 | Yes | Yes |
| **FREEMAN4** | *M. corallinus* | SRR35572337 | No | Yes |
| **GAB1** | *M. caespitosa* | SRR12424423 | No | Yes |
| **GAB2** | *M. caespitosa* | SRR12424422 | No | Yes |
| **GUT5** | Northern *M. guttatus* | SRR13618756 | No | Yes |
| **HACK** | Northern *M. decorus* | SRR10194634 | No | Yes |
| **HEC2** | Coastal *M. guttatus* | SRR35572336 | Yes | Yes |
| **HELIPORT1** | Southern *M. guttatus* | SRR35572335 | No | Yes |
| **HJA2** | Northern *M. decorus* | SRR10194632 | No | Yes |
| **HMC1** | *M. corallinus* | SRR35572334 | No | Yes |
| **HOC2** | Coastal *M. guttatus* | SRR35572333 | Yes | Yes |
| **HRH10** | Northern *M. guttatus* | SRR23709141 | No | Yes |
| **HRH16** | *M. glaucescens* | SRR23709143 | No | Yes |
| **HRL21** | *M. glaucescens* | SRR23709142 | No | Yes |
| **HUNT1** | *M. corallinus* | SRR35572332 | No | Yes |
| **HWY15D** | Northern *M. decorus* | SRR10194631 | Yes | Yes |
| **HWY70_3** | Sierran *M. guttatus* | SRR35572331 | Yes | Yes |
| **HWY70-9** | Sierran *M. guttatus* | SRR35572329 | Yes | Yes |
| **IM62** | Northern *M. guttatus* | SRR7285202 | Yes | Yes |
| **IM767** | Northern *M. guttatus* | SRX487581 | Yes | No |
| **IM851** | Northern *M. guttatus* | SRX4188698 | Yes | No |
| **IMPIA** | Northern *M. decorus* | SRR4345074 | No | Yes |
| **IMPO** | Northern *M. decorus* | SRR10194630 | Yes | Yes |
| **INV** | Southern *M. guttatus* | SRR10194629 | No | Yes |
| **KCK1** | *M. caespitosa* | SRR12424416 | No | Yes |
| **KINK** | Northern *M. decorus* | SRR10194643 | No | Yes |
| **KOOT** | *M. nasutus* | SRR1259272 | No | Yes |
| **KP1** | *M. corallinus* | SRR35572328 | No | Yes |
| **LM13** | *M. corallinus* | SRR35572327 | No | Yes |
| **LMC24** | Southern *M. guttatus* | SRR072031 | No | Yes |
| **LPD37** | Northern *M. guttatus* | SRR16646653 | No | Yes |
| **MAR3** | Northern *M. guttatus* | SRR3103524 | No | Yes |
| **Maude3** | Northern *M. decorus* | SRR35572326 | No | Yes |
| **MED84** | Sierran *M. guttatus* | SRR1298376 | Yes | Yes |
| **MEX1** | Southern *M. guttatus* | SRR35572325 | No | Yes |
| **MTC1** | Northern *M. guttatus* | SRR16646651 | No | Yes |
| **NF48.1** | Northern M. guttatus | NA | Yes | No |
| **NHN26** | *M. nasutus* | SRR1259274 | No | Yes |
| **NOR511** | *M. minor* | SRR12424414 | No | Yes |
| **NOR523** | *M. minor* | SRR12424415 | No | Yes |

| | | | | |
|---|---|---|---|---|
| **NTL3** | Northern *M. guttatus* | SRR35572324 | No | Yes |
| **OD11** | Southern *M. decorus* | SRR10194639 | Yes | Yes |
| **OD15** | Southern *M. decorus* | SRR35572323 | No | Yes |
| **OD4** | Southern *M. decorus* | SRR35572322 | Yes | Yes |
| **ODW11** | Southern *M. decorus* | SRR35572321 | No | Yes |
| **ODW13** | Southern *M. decorus* | SRR35572320 | No | Yes |
| **ODW26** | Southern *M. decorus* | SRR35572318 | No | Yes |
| **ODW36** | Southern *M. decorus* | SRR35572317 | No | Yes |
| **ODWD1** | Southern *M. decorus* | SRR35572316 | No | Yes |
| **ODWD11** | Southern *M. decorus* | SRR35572315 | No | Yes |
| **ODWD4** | Southern *M. decorus* | SRR35572314 | No | Yes |
| **ODWD6** | Southern *M. decorus* | SRR35572313 | No | Yes |
| **ODWD7** | Southern *M. decorus* | SRR35572312 | No | Yes |
| **OPB26** | Coastal *M. guttatus* | SRR35572311 | Yes | Yes |
| **OPN1** | *M. laciniatus* | SRR23709136 | No | Yes |
| **OPR105** | Sierran *M. guttatus* | SRR35572310 | Yes | Yes |
| **OSW4** | Coastal *M. guttatus* | SRR35572309 | Yes | Yes |
| **PAG2** | *M. caespitosa* | SRR12424413 | No | Yes |
| **PED5** | Southern *M. guttatus* | SRR071969 | No | Yes |
| **Per** | *M. laciniatus* | SRR23709135 | No | Yes |
| **PL20** | *M. glaucescens* | SRR35572307 | No | Yes |
| **PL22** | *M. glaucescens* | SRR35572306 | Yes | Yes |
| **PL5** | *M. glaucescens* | SRR35572305 | No | Yes |
| **PL7** | *M. glaucescens* | SRR35572304 | No | Yes |
| **PlumCk17** | *M. nasutus* | SRR35572303 | No | Yes |
| **PLUMII53** | *M. nasutus* | SRR35572302 | No | Yes |
| **PMC1** | *M. corallinus* | SRR35572301 | No | Yes |
| **PMO13** | *M. glaucescens* | SRR35572300 | No | Yes |
| **PMO3** | *M. glaucescens* | SRR23709139 | Yes | Yes |
| **PMO5** | *M. glaucescens* | SRR35572299 | No | Yes |
| **PMO8** | *M. glaucescens* | SRR35572298 | No | Yes |
| **Porch** | *M. glabratus* | SRR35572296 | No | Yes |
| **PUL18** | *M. glaucescens* | SRR35572295 | No | Yes |
| **PUL2** | *M. glaucescens* | SRR35572294 | No | Yes |
| **PUL5** | *M. glaucescens* | SRR23709138 | No | Yes |
| **PUR11** | *M. glaucescens* | SRR35572293 | No | Yes |
| **PUR4** | *M. glaucescens* | SRR35572292 | No | Yes |
| **QRY5** | Northern *M. guttatus* | SRR16646641 | No | Yes |
| **RANCHERA2** | *M. corallinus* | SRR35572291 | No | Yes |
| **ratsn7gl** | *M. glaucescens* | SRR35572290 | No | Yes |
| **rattles6** | *M. glaucescens* | SRR35572289 | No | Yes |
| **REM8** | Southern *M. guttatus* | SRR071971 | No | Yes |
| **Rogue1** | Northern *M. decorus* | SRR35572288 | Yes | Yes |

| | | | | |
|---|---|---|---|---|
| **SEQUOIA2** | Southern *M. guttatus* | SRR35572287 | No | Yes |
| **SF5** | *M. nasutus* | SRR400478 | No | Yes |
| **SHAVER3** | Southern *M. guttatus* | SRR35572285 | No | Yes |
| **SHERMAN1** | Southern *M. guttatus* | SRR35572284 | No | Yes |
| **SHG** | Southern *M. guttatus* | SRR13618755 | No | Yes |
| **SILC** | *M. corallinus* | SRR13618752 | Yes | Yes |
| **SILF** | *M. corallinus* | SRR13618751 | Yes | Yes |
| **SLP** | Southern *M. guttatus* | SRR13618751 | No | Yes |
| **SOC3** | Northern *M. guttatus* | SRR35572283 | Yes | Yes |
| **Soda1** | Southern *M. decorus* | SRR35572282 | No | Yes |
| **IM777** | Northern *M. guttatus* | SRR4345062 | No | Yes |
| **IM742** | Northern *M. guttatus* | SRR4345061 | No | Yes |
| **IM785** | Northern *M. guttatus* | SRR4345063 | No | Yes |
| **IM709** | Northern *M. guttatus* | SRR4345060 | No | Yes |
| **IM667** | Northern *M. guttatus* | SRR4345059 | No | Yes |
| **IM179** | Northern *M. guttatus* | SRR4345078 | No | Yes |
| **IM274** | Northern *M. guttatus* | SRR4345047 | No | Yes |
| **IM275** | Northern *M. guttatus* | SRR4345048 | No | Yes |
| **IM294** | Northern *M. guttatus* | SRR4345049 | No | Yes |
| **IM646** | Northern *M. guttatus* | SRR4345056 | No | Yes |
| **IM666** | Northern *M. guttatus* | SRR4345058 | No | Yes |
| **IM221** | Northern *M. guttatus* | SRR4345080 | No | Yes |
| **IM1054** | Northern *M. guttatus* | SRR4345069 | No | Yes |
| **IM909** | Northern *M. guttatus* | SRR4345067 | No | Yes |
| **SWB** | Coastal *M. guttatus* | SRR072030 | No | Yes |
| **SWB35** | Coastal *M. guttatus* | SRR35572281 | Yes | Yes |
| **Table6** | Sierran *M. guttatus* | SRR35572280 | Yes | Yes |
| **TFP17** | Sierran *M. guttatus* | SRR35572279 | No | Yes |
| **TFP3** | Sierran *M. guttatus* | SRR35572278 | No | Yes |
| **TFP6** | Sierran *M. guttatus* | SRR35572277 | Yes | Yes |
| **TFP7** | Sierran *M. guttatus* | SRR35572276 | No | Yes |
| **TMB1-14** | Sierran *M. guttatus* | SRR35572274 | No | Yes |
| **TMB2-17** | Sierran *M. guttatus* | SRR35572273 | No | Yes |
| **TMB2-22** | Sierran *M. guttatus* | SRR35572272 | No | Yes |
| **TMBI3** | Sierran *M. guttatus* | SRR35572271 | Yes | Yes |
| **TOK1** | Northern *M. guttatus* | SRR35572270 | No | Yes |
| **TOULUMNE2** | *M. corallinus* | SRR35572269 | No | Yes |
| **Trapper7** | Southern *M. decorus* | NA | Yes | No |
| **Trapper8** | Southern *M. decorus* | SRR35572268 | No | Yes |
| **TRT4** | *M. laciniatus* | SRR23709134 | No | Yes |
| **TWN36** | *M. caespitosa* | SRR12424421 | No | Yes |
| **UNP12** | *M. minor* | SRR12424420 | No | Yes |
| **UpDes24** | Northern *M. guttatus* | SRR12424420 | No | Yes |

| | | | | |
|---|---|---|---|---|
| **UTC1** | *M. caespitosa* | SRR12424419 | No | Yes |
| **UTC2** | *M. caespitosa* | SRR12424418 | No | Yes |
| **WCC1** | *M. corallinus* | SRR35572267 | No | Yes |
| **WEN4** | Northern *M. guttatus* | SRR35572266 | Yes | Yes |
| **YJS6** | Northern *M. guttatus* | SRR071970 | No | Yes |
| **YV06** | Southern *M. guttatus* | SRR10194637 | No | Yes |

**Table S2. Genes in top outlier windows (highlighted in Figure 5A).**

| Chr | Window Start | End | Name (v3.1) | Gene Start | Gene End | Alias (v2.0) |
|---|---|---|---|---|---|---|
| **Chr_02** | 75883 | 109612 | Migut.02G001000 | 73244 | 77202 | Migut.B00010.v2.0 |
| **Chr_02** | 75883 | 109612 | Migut.02G001100 | 77008 | 81685 | Migut.B00011.v2.0 |
| **Chr_02** | 75883 | 109612 | Migut.02G001200 | 86168 | 89942 | Migut.B00012.v2.0 |
| **Chr_02** | 75883 | 109612 | Migut.02G001300 | 90448 | 95316 | Migut.B00013.v2.0 |
| **Chr_02** | 75883 | 109612 | Migut.02G001400 | 95326 | 98422 | Migut.B00014.v2.0 |
| **Chr_02** | 75883 | 109612 | Migut.02G001500 | 98600 | 101310 | Migut.B00015.v2.0 |
| **Chr_02** | 75883 | 109612 | Migut.02G001600 | 101359 | 103079 | Migut.B00016.v2.0 |
| **Chr_02** | 75883 | 109612 | Migut.02G001700 | 105780 | 107737 | Migut.B00017.v2.0 |
| **Chr_02** | 75883 | 109612 | Migut.02G001800 | 107719 | 111685 | Migut.B00018.v2.0 |
| **Chr_06** | 758478 | 776286 | Migut.06G017500 | 755381 | 759686 | Migut.F00173.v2.0 |
| **Chr_06** | 758478 | 776286 | Migut.06G017600 | 759083 | 761506 | Migut.F00174.v2.0 |
| **Chr_06** | 758478 | 776286 | Migut.06G017700 | 762703 | 766717 | Migut.F00175.v2.0 |
| **Chr_06** | 758478 | 776286 | Migut.06G017800 | 766680 | 768139 | Migut.F00176.v2.0 |
| **Chr_06** | 758478 | 776286 | Migut.06G017900 | 768430 | 780568 | Migut.F00177.v2.0 |
| **Chr_06** | 4393161 | 4407635 | Migut.06G090700 | 4398844 | 4402107 | Migut.F00927.v2.0 |
| **Chr_06** | 4393161 | 4407635 | Migut.06G090800 | 4403935 | 4405503 | Migut.F00928.v2.0 |
| **Chr_06** | 4393161 | 4407635 | Migut.06G090900 | 4405639 | 4408806 | Migut.F00929.v2.0 |
| **Chr_06** | 18360349 | 18378089 | Migut.06G164300 | 18362528 | 18366931 | Migut.F01713.v2.0 |
| **Chr_06** | 18360349 | 18378089 | Migut.06G164400 | 18370815 | 18372903 | Migut.F01715.v2.0 |
| **Chr_06** | 18360349 | 18378089 | Migut.06G164500 | 18373754 | 18377252 | Migut.F01716.v2.0 |
| **Chr_08** | 21743771 | 21762253 | Migut.08G189500 | 21745841 | 21750502 | Migut.H01976.v2.0 |
| **Chr_08** | 21743771 | 21762253 | Migut.08G189600 | 21749878 | 21753885 | NA |
| **Chr_08** | 21743771 | 21762253 | Migut.08G189700 | 21755624 | 21758483 | NA |
| **Chr_08** | 21743771 | 21762253 | Migut.08G189800 | 21759462 | 21764729 | Migut.H01979.v2.0 |
| **Chr_08** | 21985394 | 22028132 | Migut.08G193200 | 22014199 | 22018274 | NA |
| **Chr_08** | 21985394 | 22028132 | Migut.08G193300 | 22018425 | 22019466 | Migut.H02015.v2.0 |
| **Chr_08** | 21985394 | 22028132 | Migut.08G193400 | 22023274 | 22026981 | Migut.H02016.v2.0 |
| **Chr_08** | 21985394 | 22028132 | Migut.08G193500 | 22027982 | 22031304 | Migut.H02017.v2.0 |
| **Chr_10** | 248068 | 270100 | Migut.10G005000 | 247610 | 249250 | Migut.J00050.v2.0 |
| **Chr_10** | 248068 | 270100 | Migut.10G005100 | 249252 | 252938 | Migut.J00051.v2.0 |
| **Chr_10** | 248068 | 270100 | Migut.10G005200 | 253233 | 256459 | Migut.J00052.v2.0 |
| **Chr_10** | 248068 | 270100 | Migut.10G005300 | 257529 | 260793 | Migut.J00053.v2.0 |
| **Chr_10** | 248068 | 270100 | Migut.10G005400 | 264171 | 267150 | Migut.J00054.v2.0 |
| **Chr_10** | 248068 | 270100 | Migut.10G005500 | 268750 | 274362 | Migut.J00055.v2.0 |
| **Chr_10** | 7927778 | 7962211 | Migut.10G103200 | 7920140 | 7944737 | Migut.J01108.v2.0 |
| **Chr_13** | 26089748 | 26145646 | Migut.13G095400 | 26102104 | 26104136 | Migut.M01025.v2.0 |
| **Chr_13** | 26089748 | 26145646 | Migut.13G095500 | 26104926 | 26106297 | Migut.M01026.v2.0 |
| **Chr_13** | 26089748 | 26145646 | Migut.13G095600 | 26106375 | 26108038 | Migut.M01027.v2.0 |
| **Chr_13** | 26089748 | 26145646 | Migut.13G095700 | 26111479 | 26112258 | Migut.M01028.v2.0 |
| **Chr_13** | 26089748 | 26145646 | Migut.13G095800 | 26112925 | 26117668 | Migut.M01029.v2.0 |
| **Chr_13** | 26089748 | 26145646 | Migut.13G095900 | 26120752 | 26124315 | Migut.M01030.v2.0 |
| **Chr_13** | 26089748 | 26145646 | Migut.13G096000 | 26126573 | 26129661 | Migut.M01031.v2.0 |
| **Chr_13** | 26089748 | 26145646 | Migut.13G096100 | 26132380 | 26133384 | NA |
| **Chr_13** | 26089748 | 26145646 | Migut.13G096200 | 26133867 | 26134382 | Migut.M01033.v2.0 |
| **Chr_13** | 26089748 | 26145646 | Migut.13G096300 | 26137461 | 26144085 | Migut.M01034.v2.0 |

| | | | | | | |
|---|---|---|---|---|---|---|
| **Chr_13** | 26089748 | 26145646 | Migut.13G096400 | 26143756 | 26145787 | Migut.M01035.v2.0 |
| **Chr_13** | 29559189 | 29576209 | Migut.13G168200 | 29558570 | 29560754 | Migut.M01753.v2.0 |
| **Chr_13** | 29559189 | 29576209 | Migut.13G168300 | 29560943 | 29561434 | Migut.M01754.v2.0 |
| **Chr_13** | 29559189 | 29576209 | Migut.13G168400 | 29565160 | 29573570 | Migut.M01755.v2.0 |
| **Chr_13** | 29559189 | 29576209 | Migut.13G168500 | 29573130 | 29575729 | Migut.M01756.v2.0 |
| **Chr_13** | 29559189 | 29576209 | Migut.13G168600 | 29575868 | 29579830 | Migut.M01757.v2.0 |
| **Chr_14** | 25922119 | 25965758 | Migut.14G269700 | 25918625 | 25940135 | Migut.N02861.v2.0 |
| **Chr_14** | 25922119 | 25965758 | Migut.14G269800 | 25943445 | 25965884 | Migut.N02862.v2.0 |
| **Chr_14** | 27365674 | 27385459 | Migut.14G299000 | 27368803 | 27370867 | Migut.N03151.v2.0 |
| **Chr_14** | 27365674 | 27385459 | Migut.14G299100 | 27371112 | 27374461 | Migut.N03152.v2.0 |
| **Chr_14** | 27365674 | 27385459 | Migut.14G299200 | 27374460 | 27376315 | Migut.N03153.v2.0 |
| **Chr_14** | 27365674 | 27385459 | Migut.14G299300 | 27377699 | 27380825 | Migut.N03154.v2.0 |
| **Chr_14** | 27365674 | 27385459 | Migut.14G299400 | 27381472 | 27384114 | Migut.N03155.v2.0 |
| **Chr_14** | 27365674 | 27385459 | Migut.14G299500 | 27384507 | 27386776 | Migut.N03156.v2.0 |

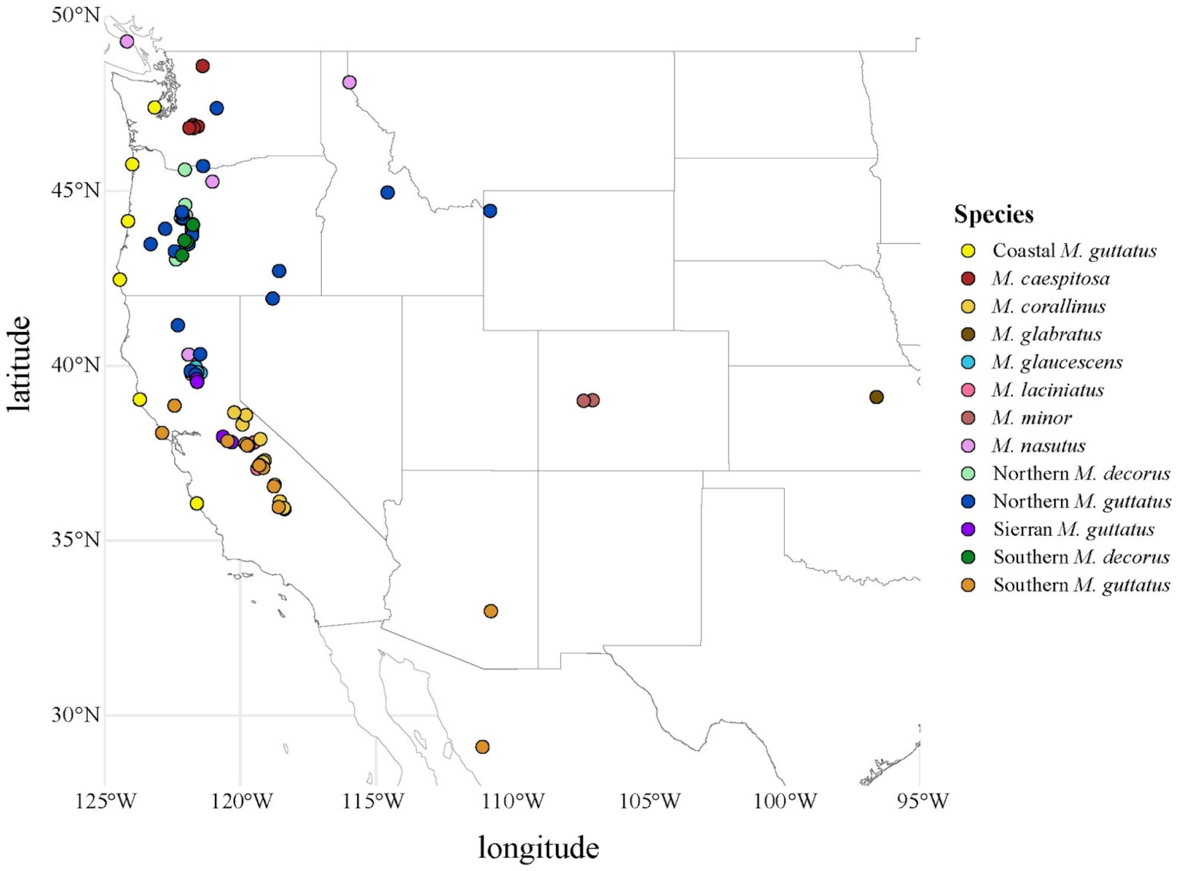


**Figure S1.** The geographic distribution of samples used for genomic analyses. Samples are colored according to species/lineage.

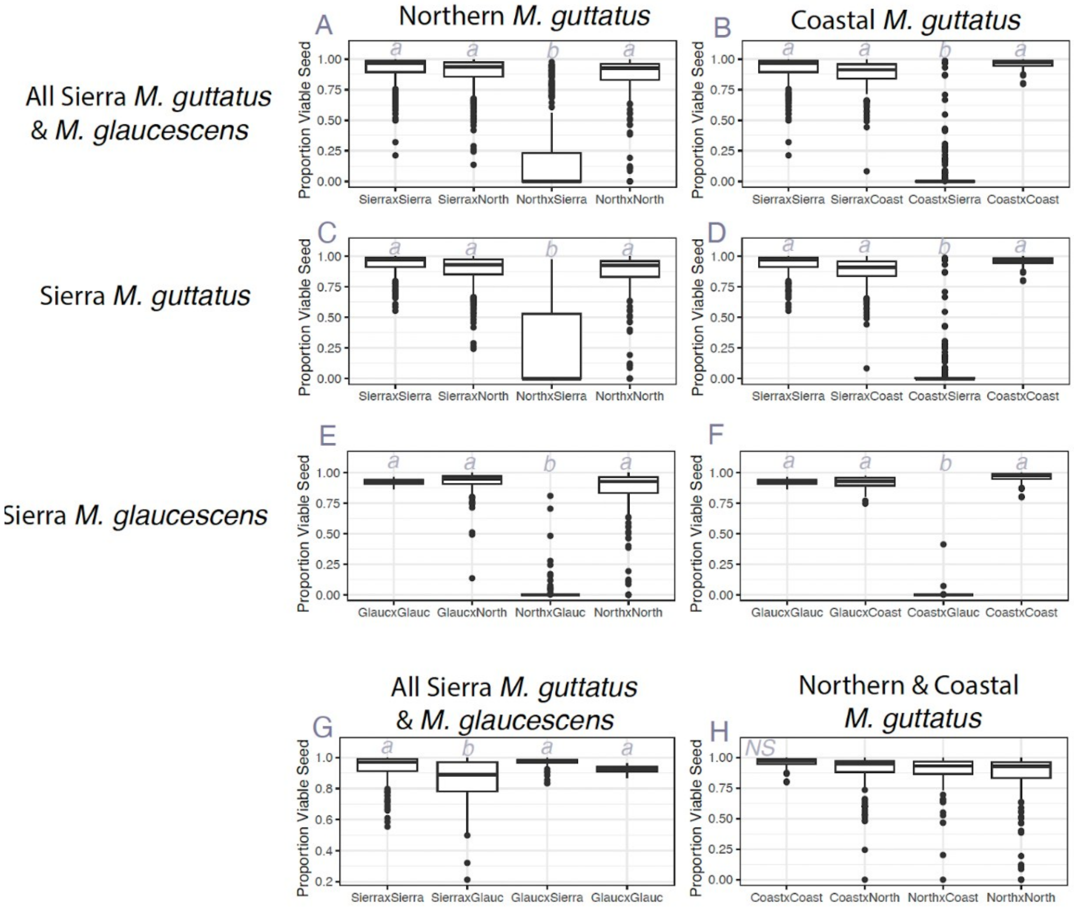


**Figure S2:** Proportion of viable seeds for lineage groups in Figure 1. (A,B) all Sierran samples (including Sierran *M. guttatus* and *M. glaucescens*). (C,D) Only Sierran *M. guttatus*. (E,F) Only Sierran *M. glaucescens*. Crosses involving Northern *M. guttatus* are on the left, and Coastal *M. guttatus* are on the right. (G,H) Crosses within groups: All Sierran lineages (G) and Northern and Coastal *M. guttatus* (H).

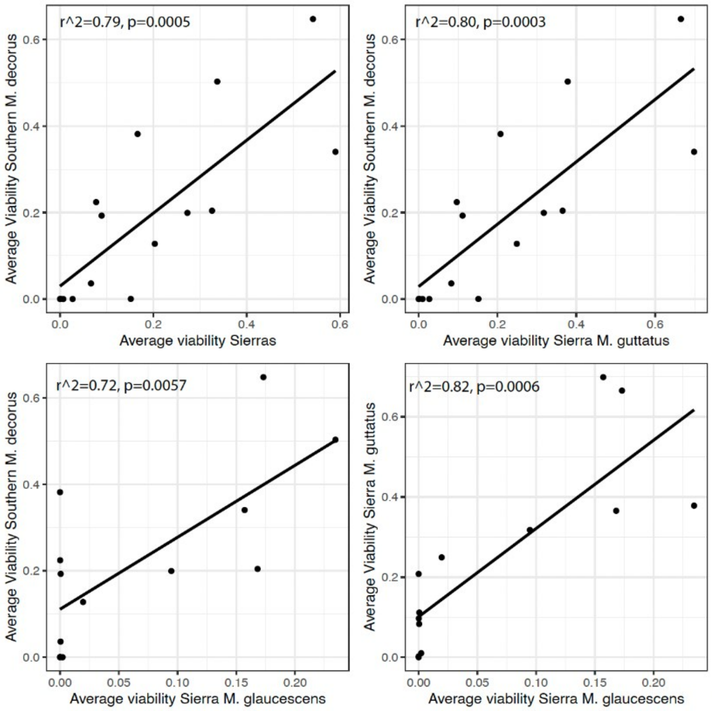


**Figure S3.** Correlations among Southern *M. decorus* and different groupings of Sierran lineages for their crossabilities (as the sire) to lines of *M. guttatus*. (A) Southern *M. decorus* and all Sierran lineages. (B) Southern *M. decorus* and only Sierran *M. guttatus*. (C) Southern *M. decorus* and only Sierran *M. glaucescens*. (D) Sierran *M. guttatus* and Sierran *M. glaucescens*. In all panels each point is the average seed viability for a line of Northern or Coastal *M. guttatus*.

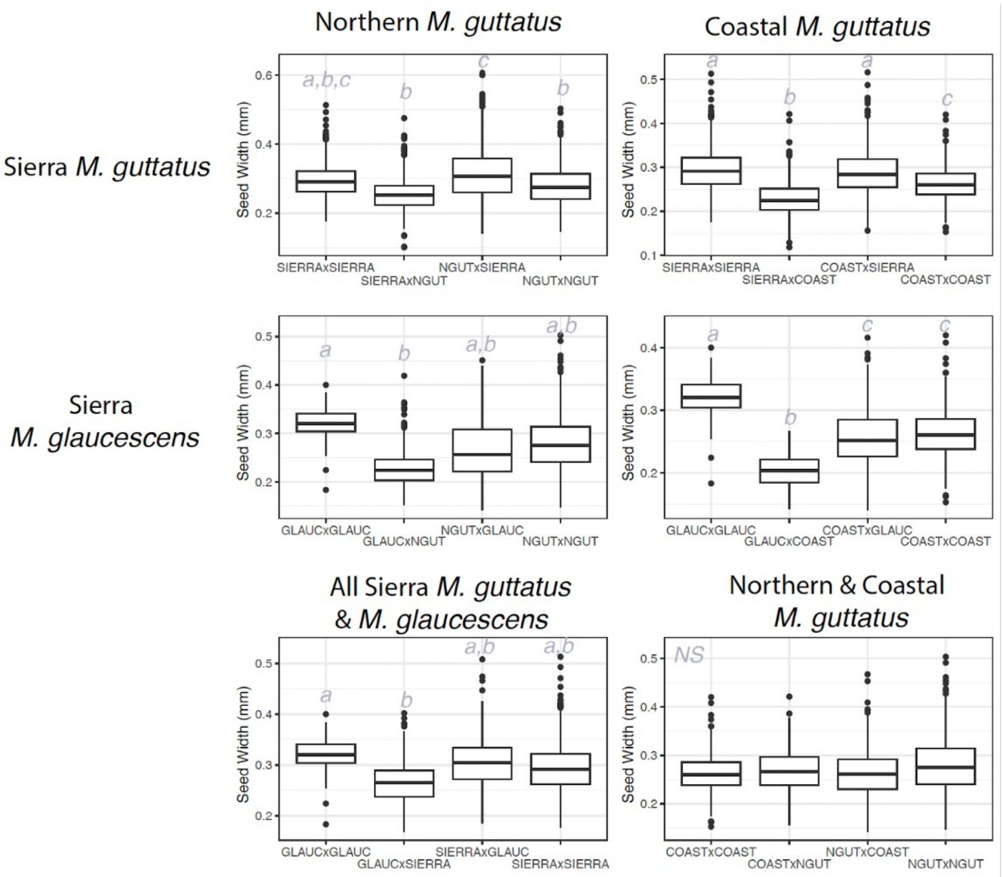


**Figure S4.** Seed sizes for the range-wide crossing survey by cross type.

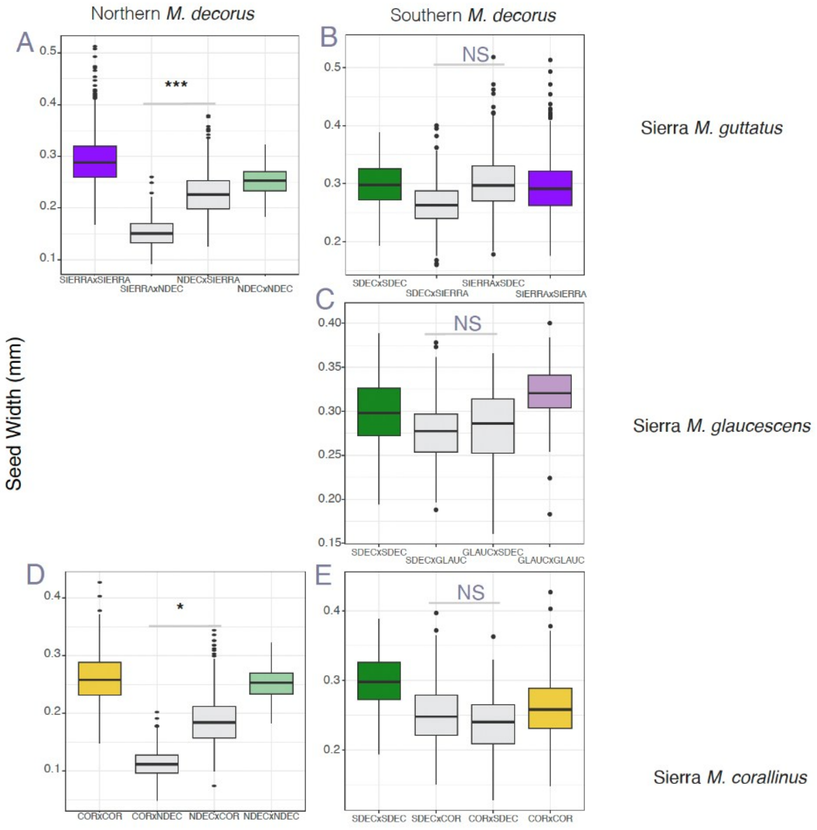


**Figure S5.** Seed sizes for crosses between Sierran lineages (*M. guttatus, M. glaucescens,* and *M. corallinus*) and the two *M. decorus* lineages with different histories of conflict.

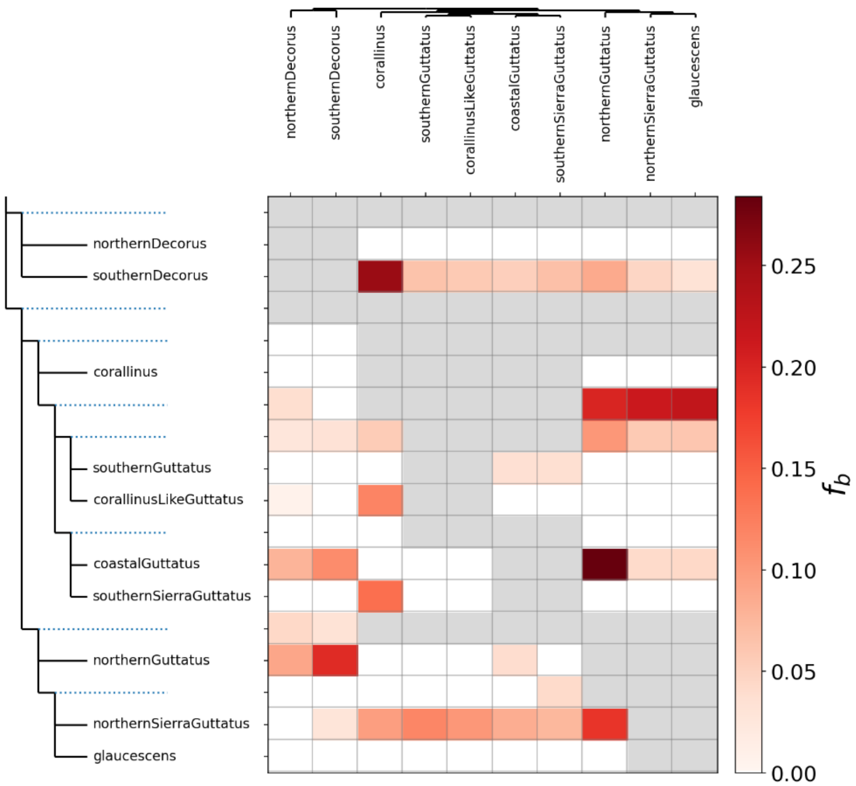


**Figure S6.** Heatmap of the f-branch statistic for select members of the *M. guttatus* species complex with varying patterns of hybrid seed inviability with each other. In all cases, *M. caespitosa* is used as the outgroup.

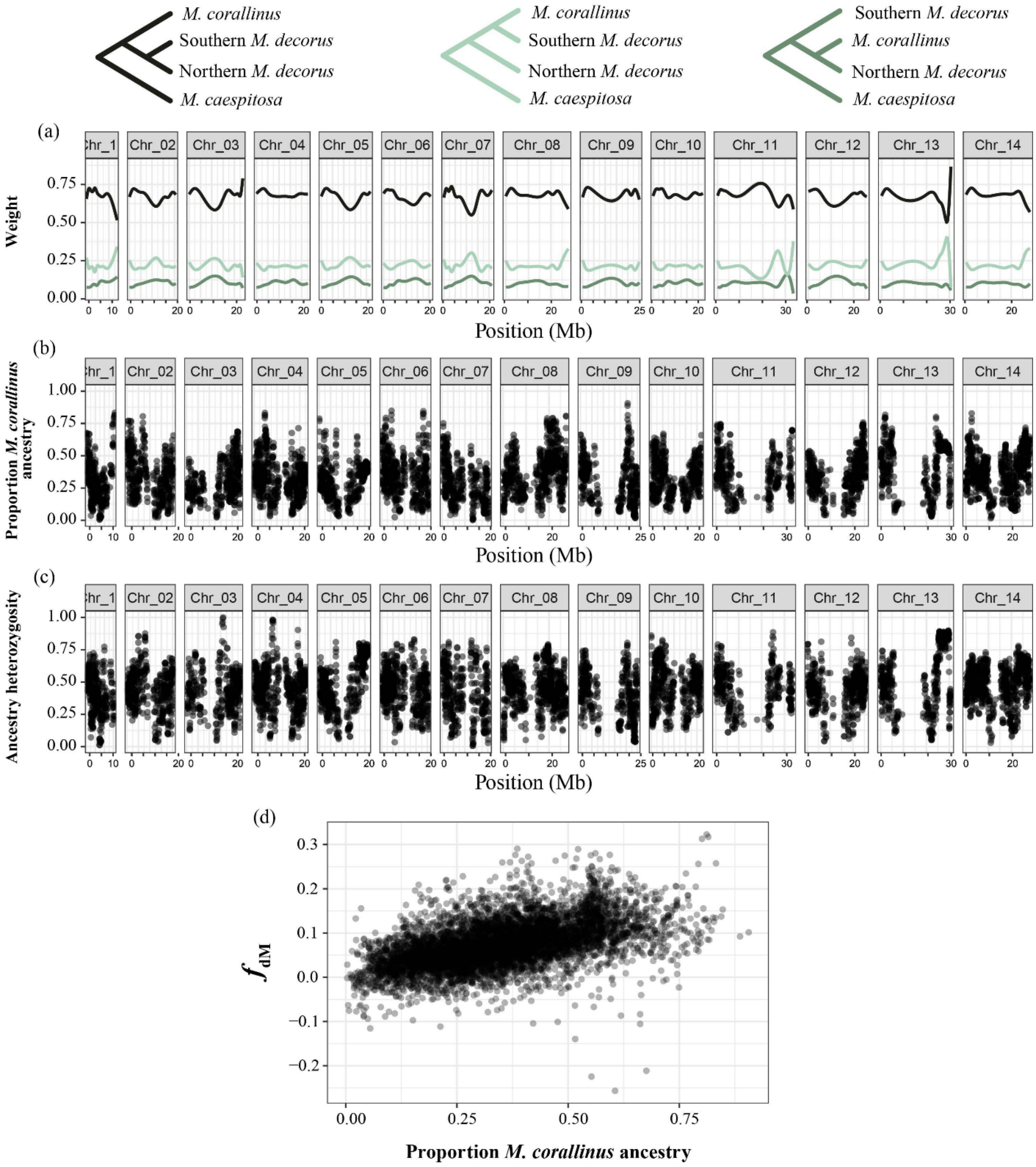


**Figure S7:** TWISST and local ancestry results across the chromosomes. A) Topology weighting was calculated for the three topologies shown in the key, in sliding windows containing 250 SNPs with a 50 SNP step between windows. Results depicted are smoothed results across the genome. B) Mean proportion of *M. corallinus* ancestry in sliding windows across the genomes of Southern *M. decorus*, estimated with AncestryHMM. C) Mean proportion of heterozygous ancestry in sliding windows across the genomes of Southern *M. decorus*, estimated with AncestryHMM. D) Relationship between the proportion of *M. corallinus* ancestry and $f_{dM}$ in sliding windows.

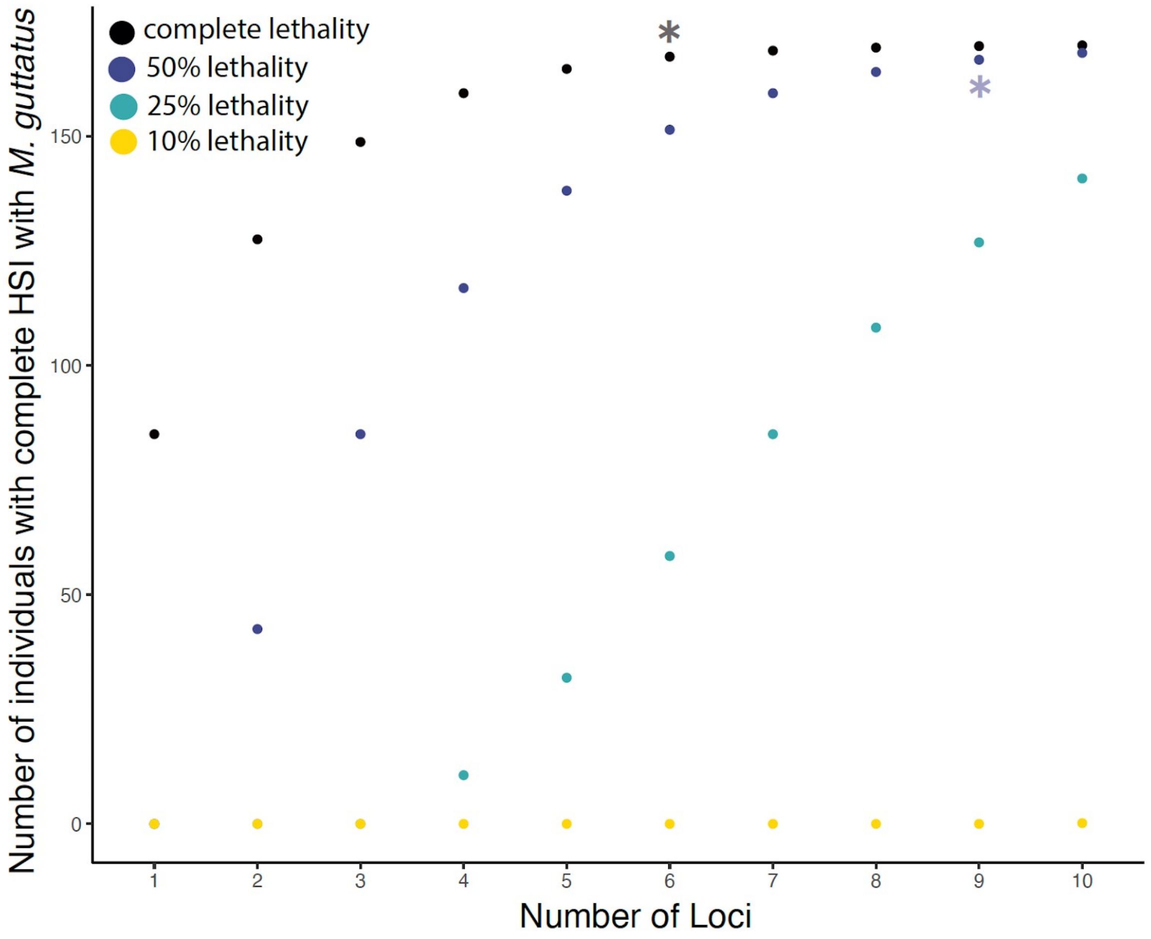


**Figure S8.** Simulations for power analysis of complementation test. We assessed how many F2 individuals (out of the 170 F2s tested) were predicted to have complete hybrid seed inviability when acting as the sire in crosses with Northern *M. guttatus* as a function of the number of loci involved in the incompatibility and their effect size. We then ran Fisher's Exact tests on the observed versus expected ratios of fully incompatible individuals to individuals with any amount of compatibility to assess whether we could distinguish having no phenotypic segregation among the F2s versus not having the power to identify individuals with any level of compatibility. For the most conservative model (i.e., each locus confers complete lethality), we could not distinguish observed vs expected results for models of 6 loci or greater. For the second most conservative model (i.e., each locus confers 50% inviability), we could not distinguish expected versus observed results for models containing 9 loci or greater. We note that in the limited studies to date of the genetic basis of hybrid seed inviability, empirical work typically implicates 3-6 paternally acting loci with effect sizes ranging from 20-50% lethality.

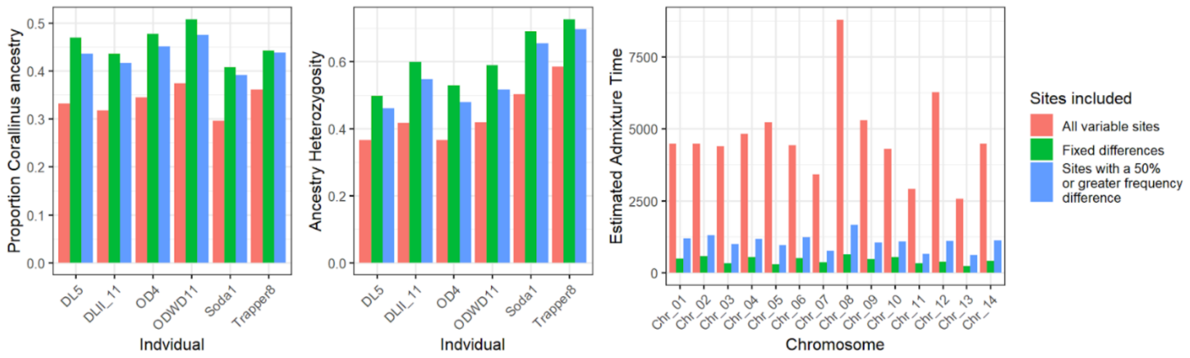


**Fig.S9:** Comparison of the inference of ancestry when using only loci that are ancestry informative between Northern *M. decorus* and *M. corallinus*. A) Proportion of corallinus ancestry in each individual. B) Proportion of SNPs which are heterozygous for ancestry in each individual. C) Estimated timing of admixture for each chromosome.

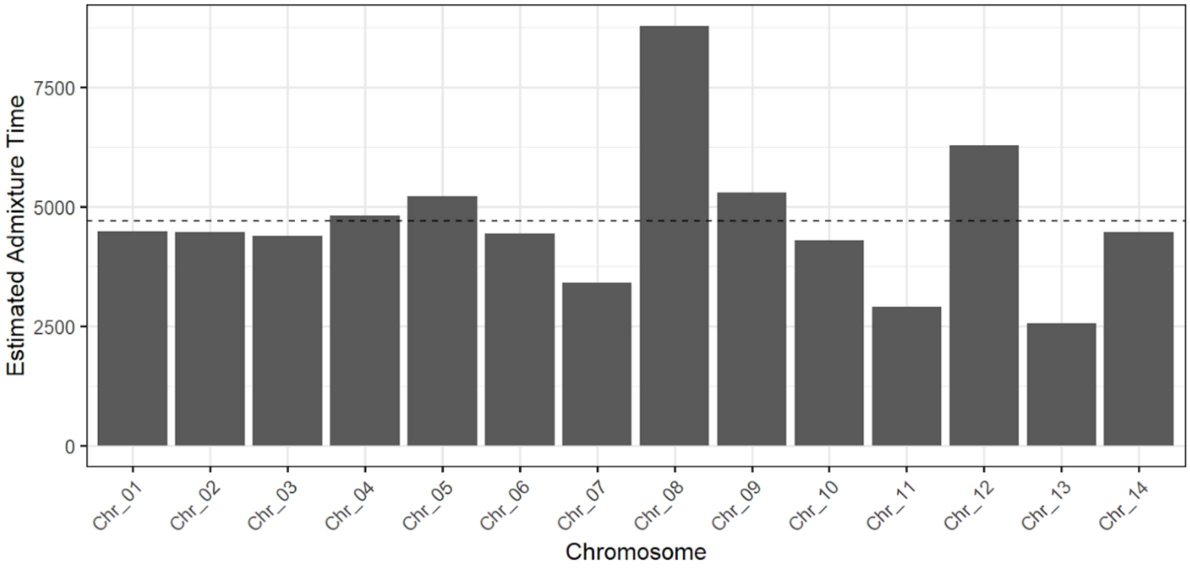


**Fig.S10:** Estimates of the timing of admixture in generations for each chromosome based on all variable sites between *M. corallinus* and Northern *M. decorus*. The dashed line indicates the average estimate (4,703 generations).
